# Serum proteomic profiling at diagnosis predicts clinical course, and need for intensification of treatment in inflammatory bowel disease

**DOI:** 10.1101/2020.08.31.276162

**Authors:** R Kalla, AT Adams, D Bergemalm, S Vatn, NA Kennedy, P Ricanek, J Lindstrom, A Ocklind, F Hjelm, NT Ventham, GT Ho, C Petren, IBD-Character Consortium, D Repsilber, J Söderholm, M Pierik, M D’Amato, F Gomollón, C Olbjorn, J Jahnsen, MH Vatn, J Halfvarson, J Satsangi

## Abstract

**Background:** Success in personalised medicine in complex disease is critically dependent on biomarker discovery. We profiled serum proteins using a novel proximity extension assay (PEA) to identify diagnostic and prognostic biomarkers in inflammatory bowel disease (IBD).

**Methods:** We conducted a prospective case-control study in an inception cohort of 552 patients (328 IBD, 224 non-IBD), profiling proteins recruited across 6 centres. Treatment escalation was characterised by the need for biological agents or surgery after initial disease remission. Nested leave-one-out cross validation was used to examine the performance of diagnostic and prognostic proteins.

**Results:** A total of 66 serum proteins differentiated IBD from symptomatic non-IBD controls including Matrix Metalloproteinase-12 (Holm adjusted p=4.1×10^−23^) and Oncostatin-M (OSM, p=3.7×10^−16^). Nine of these proteins associate with *cis*- germline variation (59 independent SNPs). Fifteen proteins, all members of TNF independent pathways including interleukin-1 and OSM predicted escalation, over a median follow-up of 518 (IQR 224-756) days. Nested cross-validation of the entire data set allows characterisation of 5-protein-models (96% comprising five core proteins ITGAV, EpCAM, IL18, SLAMF7, and IL8) which define a high-risk subgroup in IBD (HR 3.90, 95% CI: 2.43-6.26), or allows distinct 2, and 3 protein models for UC and CD respectively.

**Conclusion:** We have characterised a simple oligo-protein panel that has the potential to identify IBD from symptomatic controls and predicts the evolution of disease over time. The technology could be suitable as a point of care testing in defining risk. Further prospective work is required to characterise the utility of the approach.

## Introduction

Personalised medicine is now a major priority in healthcare research. Programmes such as the 7^th^ framework programme for research and technological development and 100,000 genomes project (www.genomicsengland.co.uk) in the UK prioritise the discovery and validation of novel biomarkers in human diseases^1^. This impetus to redefine clinical practice coupled with an increasingly wide therapeutic choice of biological agents, and small molecules has driven interest in risk-stratifying patients at diagnosis in Inflammatory Bowel Disease (IBD)^2–4^.

There have been recent scientific advances catalysing biomarker discovery studies. It is now apparent that genes that contribute to prognosis in Crohn’s disease (CD) are distinct from those that predict disease susceptibility4. Studies in both adults and children have demonstrated that patients with a progressive disease display a unique transcriptional signature^3,5–7^. Critically for translation, emergent data demonstrate that early biomarker-driven therapeutic interventions can improve disease outcomes in CD^8^.

Despite significant progress in multi-omic biomarker discoveries, none are in routine clinical use. Markers such as c-reactive protein (CRP) have shown clinical utility in disease susceptibility, activity and behaviour^1^. Faecal calprotectin (FC) however has emerged to date as the most reliable and accurate diagnostic protein biomarker in IBD9. Recently, randomised trial data demonstrate that early biomarker-driven therapeutic interventions based on FC can improve disease outcomes in CD^8^. However, there are well-described limitations of faecal testing in clinical care^2,10,11^ that highlight the need for blood-based markers to maximise uptake and acceptability.

Multiprotein signatures have potentially diverse clinical applications from early detection of IBD to disease classification and behaviour, response to therapy, and monitoring disease activity. Technological limitations in multi-protein profiling have recently been overcome^12,13^, with the discovery of innovative approaches for multiplexing biological samples utilizing minimal sample volume but providing a highly sensitive and specific immunoassay. Proximity extension assays (PEA) are antibody-based methods that utilise two or more DNA-tagged aptamers or antibodies that bind when in close proximity to the target protein or protein complex. PEA allows multiplexing with 1 microlitre (μL) sample consumption, and a high sensitivity and specificity for proteins of interest^12,13^

In this report, we explore the diagnostic and prognostic capabilities of circulating PEA based proteins markers in IBD and their association with germline variations. Our study demonstrates that protein panels can predict disease and its course.

## Materials and Methods

### Study Design

We conducted a prospective, multi-centre case-control study in patients with suspected or confirmed IBD, recruited at presentation either as in-patients or electively as out-patients across 6 clinical centres in UK and Europe (EU Character reference no. 305676). Demographic data including age, sex, date of diagnosis (**Table 1**) and details of drug therapies were collected. Treatment naivety within the IBD cohort was defined as no exposure to any IBD related medical therapies such as steroids, 5-ASA, biologics and immunomodulators (**Supplementary Table 1**). Blood samples for protein profiles and genotyping were collected at baseline at the time of recruitment. High sensitivity C-reactive protein (hsCRP), albumin, and faecal calprotectin (if stool had been collected around recruitment), were re-assayed in a single batch at the end of recruitment. Other routine markers were tested as part of routine clinical care. Clinical outcome data were collected at follow up for patients with IBD.

**Table 1A:**
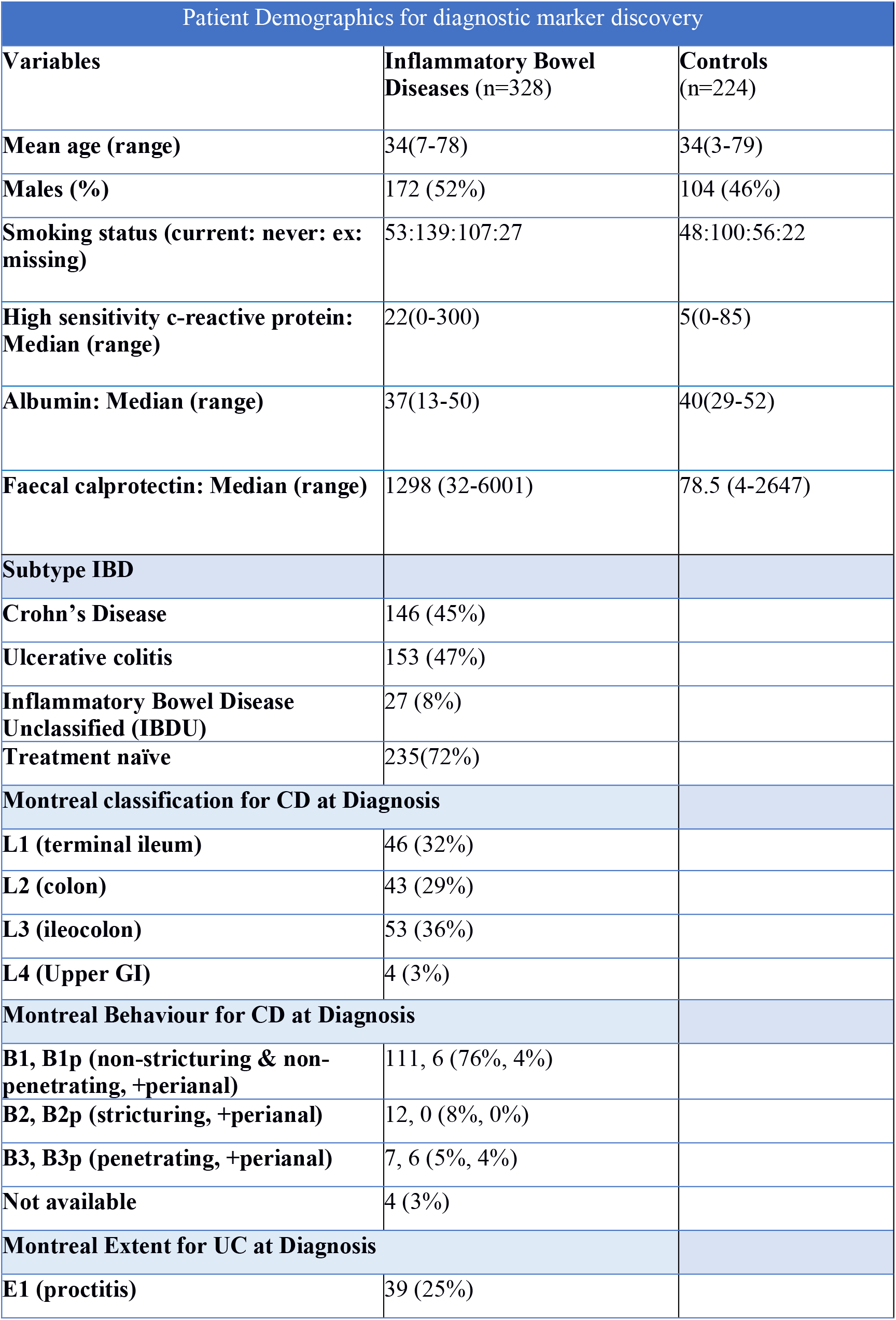
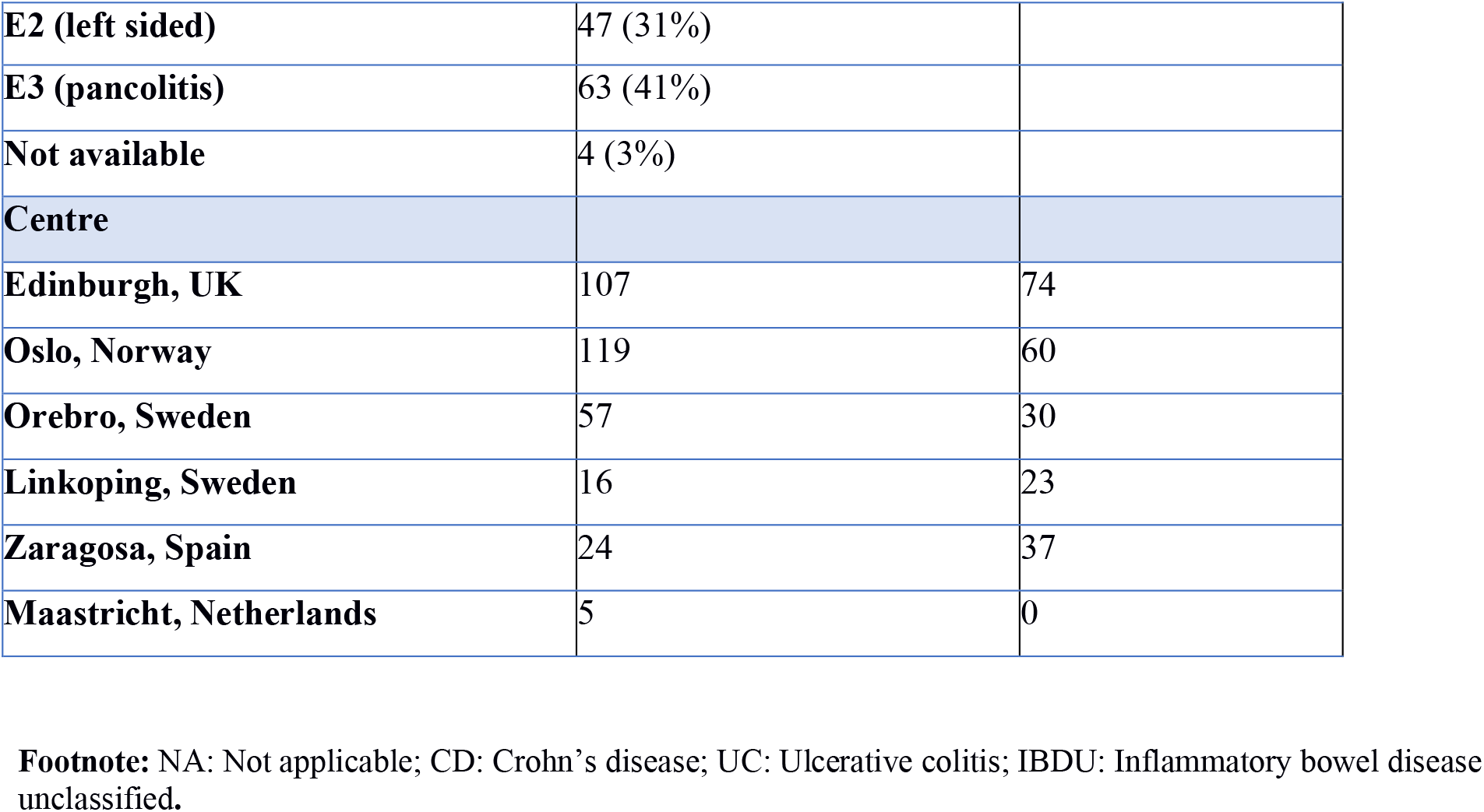
Patient demographics of patients included in our study of protein expression in newly diagnosed inflammatory bowel disease and symptomatic controls.

**Table 1B:** Patient demographics for predicting disease course in Inflammatory Bowel Disease

| Inflammatory Bowel Disease |  |  |
| --- | --- | --- |
| Variables | IBD escalation group (n=67) | Non-escalation group (n=212) |
| Males (%) | 39(58) | 109(51) |
| Smoking status (current: never: ex: missing) | 16:34:16:1 | 36:98:77:1 |
| Median FC (range) | 1631 (35-6001) | 1186 (32-6001) |
| Median age (range) | 28(18-67) | 33(18-77) |
| Edin: Norway: Sweden: Spain | 26:22:15:4 | 81:71:41:19 |
| Disease subtype |  |  |
| Crohn's disease | 39 | 73 |
| Ulcerative colitis | 26 | 117 |
| Inflammatory bowel disease unclassified (IBDU) | 2 | 22 |

| Ulcerative Colitis |  |  |
| --- | --- | --- |
| Variables | Escalation group (n=26) | Non-escalation group (n=117) |
| Males (%) | 19(73) | 67(57) |
| Smoking status (current: never: ex: missing) | 3:9:14:0 | 8:53:56:0 |
| Median FC (range) | 3778 (35-6001) | 1367 (32-6001) |
| Median age (range) | 30(18-60) | 37(18-77) |
| Edin: Norway: Sweden: Spain | 13:8:4:1 | 39:52:19:7 |
| Paris Extent for UC |  |  |
| E1 (proctitis) | 0 | 38 (32%) |
| E2 (left sided) | 7 (27%) | 37 (32%) |
| E3 (pancolitis) | 19 (73%) | 42 (36%) |

| Crohn's Disease |  |  |
| --- | --- | --- |
| Variables | Escalation group (n=39) | Non-escalation group (n=73) |
| Males (%) | 19(49) | 33(45) |
| Smoking status (current: never: ex: missing) | 13:5:20:1 | 26:18:28:1 |
| Median FC (range) | 1398.5 (47-6001) | 825 (70-6001) |
| Median age (range) | 25(18-66) | 29(18-73) |
| Edin: Norway: Sweden: Spain | 11:14:11:3 | 34:17:12:10 |
| Montreal classification for CD |  |  |
| L1 (terminal ileum) | 13 (33%) | 25 (34%) |
| L2 (colonic) | 9 (23%) | 22 (30%) |
| L3 (ileocolon) | 17 (44%) | 25 (34%) |
| L4 (upper GI) | 0 | 1 (1%) |
| Montreal Behaviour for CD |  |  |
| B1, B1p (non-stricturing & non-penetrating, +perianal) | 29, 0 (74%, 0%) | 55, 6 (75%, 8%) |
| B2, B2p (stricturing, +perianal) | 6, 0 (15%, 0%) | 4, 0 (5%, 0%) |
| B3, B3p (penetrating, +perianal) | 2, 2 (5%, 5%) | 5, 1 (7%, 1%) |
| Not available | 0 | 2 (3%) |

### Inclusion criteria

Patients with a suspected or new diagnosis of IBD were included in the study; prospectively recruited from out-patients and in-patient settings across participating centres. All IBD cases met the standard diagnostic criteria for Ulcerative colitis (UC), CD or Inflammatory Bowel Disease Unclassified (IBDU) following thorough clinical, microbiological, endoscopic, histological, and radiological evaluation. The Lennard-Jones, Montreal and Paris criteria were used for diagnosis and classification of clinical phenotypes^14–16^. The control group consisted of patients with gastrointestinal symptoms (symptomatic controls) who had no discernible evidence of IBD at any time during follow-up.

### Clinical Course in IBD

The primary end-point of treatment escalation was defined as the need for a biologic, ciclosporin or surgery, instituted for disease flare after initial induction therapy and aiming to induce disease remission. In UC, the definition of treatment escalation included any patient requiring colectomy during their index admission.

### Sample collection and processing

We collected blood samples prospectively and processed serum within two hours of sampling (Vacuette® gel tube with clot activator and using centrifugation at 2000G for 10 minutes). Serum was subsequently stored at −80°C until further use.

We measured protein concentrations using Proximity Extension Assay technology^12^. For each panel, 92 oligonucleotide-labelled antibody probe pairs are allowed to bind to their respective target present in the sample. A PCR reporter sequence is formed by a proximity-dependent DNA polymerization event and is subsequently detected and quantified using real-time PCR. Four internal controls were included in each multiplex reaction, and negative controls and an interplate control samples were included on each assay plate.

Whole-blood leukocyte DNA was extracted using the Nucleon BACC 3 DNA extraction kit (GE healthcare, Buckinghamshire, UK). We genotyped patients using the Illumina OmniExpressExome-8 Bead Chip (Illumina, San Diego, CA, USA).

### IBD Protein Panel Design

We generated a candidate list of IBD genetic risk loci using the published genome-wide association studies^17,18^ and other sources from the literature relevant to IBD biology. After thorough quality control, assay analyses and validation, we developed a strategy designed to allow the incorporation of a total of 460 commercially available protein antibodies into five novel multiplex protein panels comprising proteins involved in IBD-related mechanisms, such as inflammation, immune regulation, metabolism and cell-cell signalling (**Supplementary Table 2**). Certain panels including the Inflammatory Olink panel are now commercially available.

### Data normalisation and quality control

Raw data (qPCR Ct values) were normalized for technical variation (extension control) and variation between multiple experimental runs (inter-plate control). The data were then adjusted with a predetermined correction factor and reported as an arbitrary unit: normalized protein expression on a log_2_ scale as described previously^19^.

Limit of detection (LOD) for each protein probe was defined as the mean plus three standard deviations of the negative controls. For quality control reasons in designing assays, we excluded 147/460 proteins where >50% of samples were below the LOD and excluded 33 samples in which >20% of the remaining proteins were below the LOD.

### Statistical analysis

We used R 3.4.4 (R Foundation for Statistical Computing, Vienna, Austria) and Julia 1.1.020 for analysis. Data were corrected for centre batch effects using ComBat. P-values were adjusted for multiple testing (Holm correction)^21^. Survival analysis was performed using Cox proportional hazard models, and diagnostic analysis with binomial logistic regression. We constructed models and characterised their predictive performance using a rigorous cross-validation approach wherein feature selection and parameter estimation were performed in an inner LOO loop, with the model performance assessed using the unseen outer LOO sample. Reported performance of the models is based on the combined performance in each outer LOO sample of the models derived in their respective inner loops. Models were constrained to include age and sex, with proteins added in a forward stepwise approach based on AIC. The number of included proteins was based on the AIC evidence ratio assessed in the first 10% of outer loops after which models were constrained to the selected number of proteins to reduce computation. No pre-selection or filtering of the proteins by any criteria was used prior to the cross-validation. Classification was based on the optimum threshold from ROC analysis of the outer cross-validation loop. Randomly permuted data (n=50) were analysed with the same technique with true data outperforming every permuted dataset.

### Genotyping and Protein Quantitative trait loci analyses (pQTLs)

Genome Studio files were imported into R for sex mismatch removal, and further analysis. Protein quantitative trait loci (pQTLs) were found using the matrix eQTL package^22^ with a distance threshold of 300Kb and a MAF threshold of >0.1. Age and sex were included as covariates, and Holm correction was applied to p values. Further sub-analysis was performed with treatment exposure, sex, age, BMI, clinical centre, and smoking status as covariates.

### Ethics Statement

All centres were granted local ethics approval for this study and all patients gave written and informed consent prior to participating in this study.

## Results

### Differentially expressed protein markers in Inflammatory Bowel Diseases

We designed PEA assays for 313 IBD-related proteins and analysed these in 552 patients recruited in six IBD centres in Europe between May 2012 and September 2015 (**Table 1**). Linear models with age and sex as covariates identified a total of 66 protein markers that showed significant differential expression between IBD (n=328) and controls (n=224, **Figure 1 and Supplementary Table 3**) including Matrix Metallopeptidase-12 (MMP-12, log_2_fold change (log_2_FC) =0.87, Holm p=4.1×10^−23^) and Oncostatin-M (OSM, log_2_FC =0.81, p=3.7×10^−16^). Over-expression in IBD was more frequent at higher significance levels (p=0.01), with the top 12 proteins all being over-expressed. Of the proteins down-regulated in disease the most significant include Growth Arrest-Specific-6 (GAS6) and Integrin alpha-V (ITGAV).

**Figure 1:**
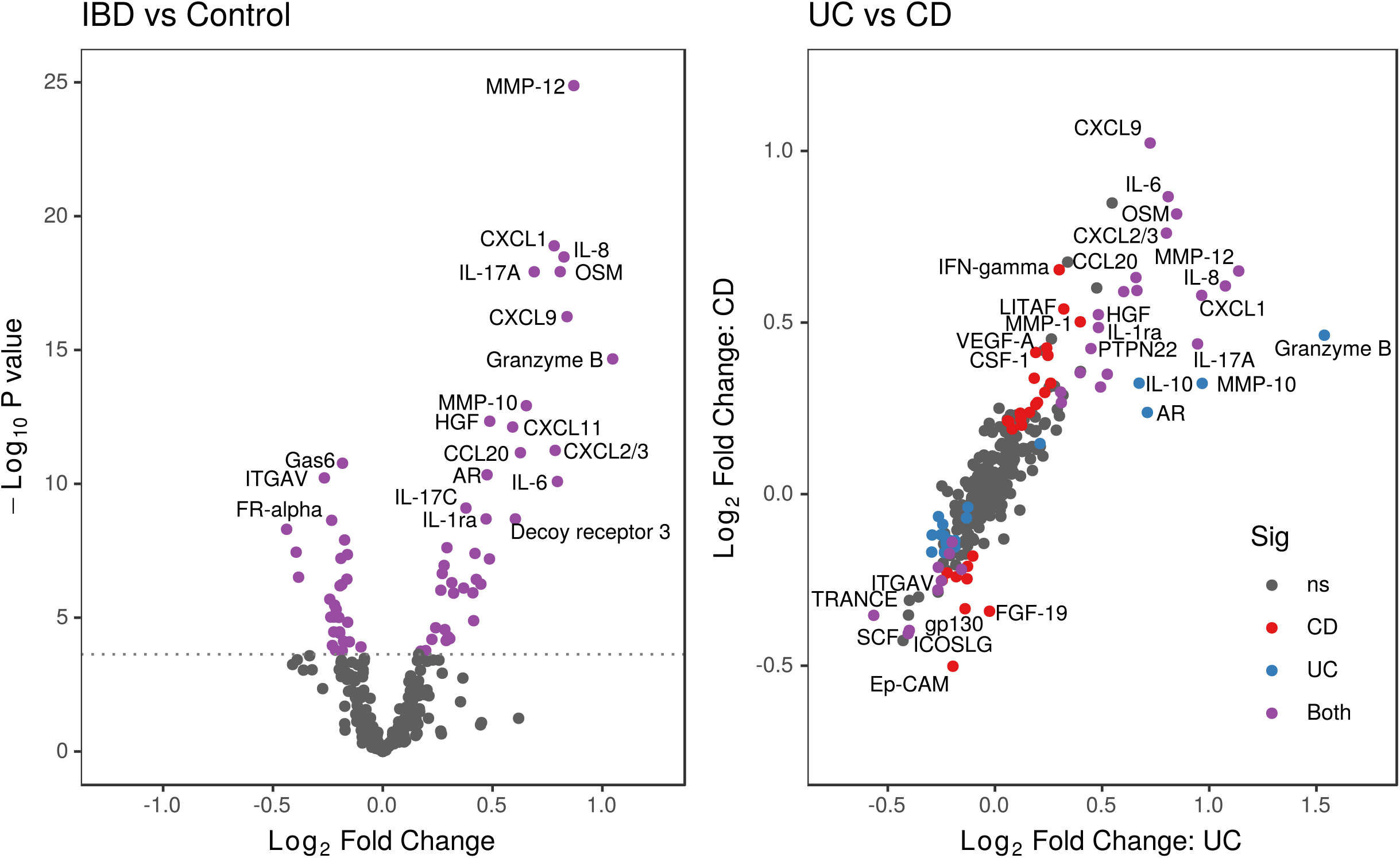
A) Volcano plot displaying the log_2_ fold-change and significance of protein associations with IBD. Dotted line indicates threshold for significance after Holm correction. B) Fold change between Ulcerative colitis(UC) and Crohn’s disease(CD) respectively vs controls, points coloured by significance after Holm correction in CD, UC, both, or neither (ns).

There were 55 protein markers that were significantly differentially expressed in CD compared to controls (**Supplementary Table 4**); the most significant being CXCL9 (log_2_FC =1.02, p=5.0×10^−15^) and OSM (log_2_FC =0.82, p=5.8×10^−12^). In ulcerative colitis (UC), 46 protein markers had significant expression differences compared to controls (**Supplementary Table 5**), including MMP-12 (log_2_FC =1.14, p=3.6×10^−26^) and Granzyme-B (log_2_FC =1.54, p=7.9×10^−23^). A total of 5 proteins showed significant expression differences between UC and CD (**Supplementary Table 6, Figure 1B**), all were significantly different between CD and controls, and differed further in the same direction in UC. A clinically useful model to distinguish between CD and UC could not be established, the best performing classifier (consisting of age, sex, and expression of six proteins) was only 68.0% accurate. Correlations between protein expression and inflammatory markers are shown in **Supplementary Figure S1.**

### Diagnosis of IBD with PEAs and inflammatory markers

We next examined the diagnostic performance of PEA-based protein models using the nested cross-validation approach, independent of the differential expression analysis. Fitting logistic regression models comprising age, sex, and 6 protein expression values in a nested cross-validation approach was 79.8% (95% CI 76.4-83.2) accurate at distinguishing IBD from controls (sensitivity 83.1%, CI 79.1-87.2; specificity 74.8%, CI 69.0-80.5). The proteins selected by each inner cross-validation loop were stable, comprising Granzyme-B (selected by 100% of inner loops), MMP12 (100%), Gas6 (99.8%), IL7 (99.6%), IL8 (99.6%), and EMMPRIN (99.3%).

This approach outperformed an hsCRP-model with age and sex, which had a sensitivity, specificity and accuracy of 77.5%(72.7-82.3), 27.8% (21.5-34.0) and 57.2% (52.9-61.7) respectively (Table 3). FC performed better (sensitivity 85.4%, CI 78.1-92.7; specificity 88.4%, CI 78.8-98.0, accuracy 86.4%, CI 80.5-92.2%), however FC suffers from poor uptake, with only 30.4% of patients having a result between 30 days prior- and 7 days post-inclusion.

The PEA-based models performed similarly in UC and CD (accuracy 78.4 & 77.7% respectively), and separate analysis of CD and UC did not produce more accurate models. FC was more sensitive in UC compared to CD (90.7, CI 83.0-98.5 vs 77.4%, CI 62.7-92.1; χ2 p=1.2×10^−12^), yielding an improved accuracy of 89.7%, CI 83.6-95.7 vs 83.8%, CI 75.4-92.2 (**Table 3**).

### Individual proteins associated with treatment escalation

In order to identify proteins that associate with treatment escalation, we analysed data from 279 patients with confirmed IBD diagnoses where follow up data were available (**Table 1B and Supplementary Table 7**). Patients who required escalation were younger (median age 28 vs 33, p=0.02), more likely to be male (58.2 vs 51.4%, χ2 p>0.05), and have CD (58.2 vs 34.4%, χ2 p=0.004). The association between treatment escalation and smoking status was not statistically significant in either CD or UC.

Cox models were created to identify protein markers individually associated with treatment escalation in IBD, accounting for age and sex. Fifteen proteins (**Figure 2 and Table 2**) were significantly associated with treatment escalation in all IBD including ITGAV (Holm p=3.2×10^−6^) and EpCAM (p=1.7×10^−4^). In UC (n=143), 22 proteins were significantly associated with treatment escalation (**Supplementary Table 8**), but in CD (n=112) no individual proteins achieved significance, although the results were correlated with those obtained for UC alone (r=0.56, p=6.6×10^−15^). Adjusting for treatment naivety did not influence the top differentially expressed proteins.

**Figure 2:**
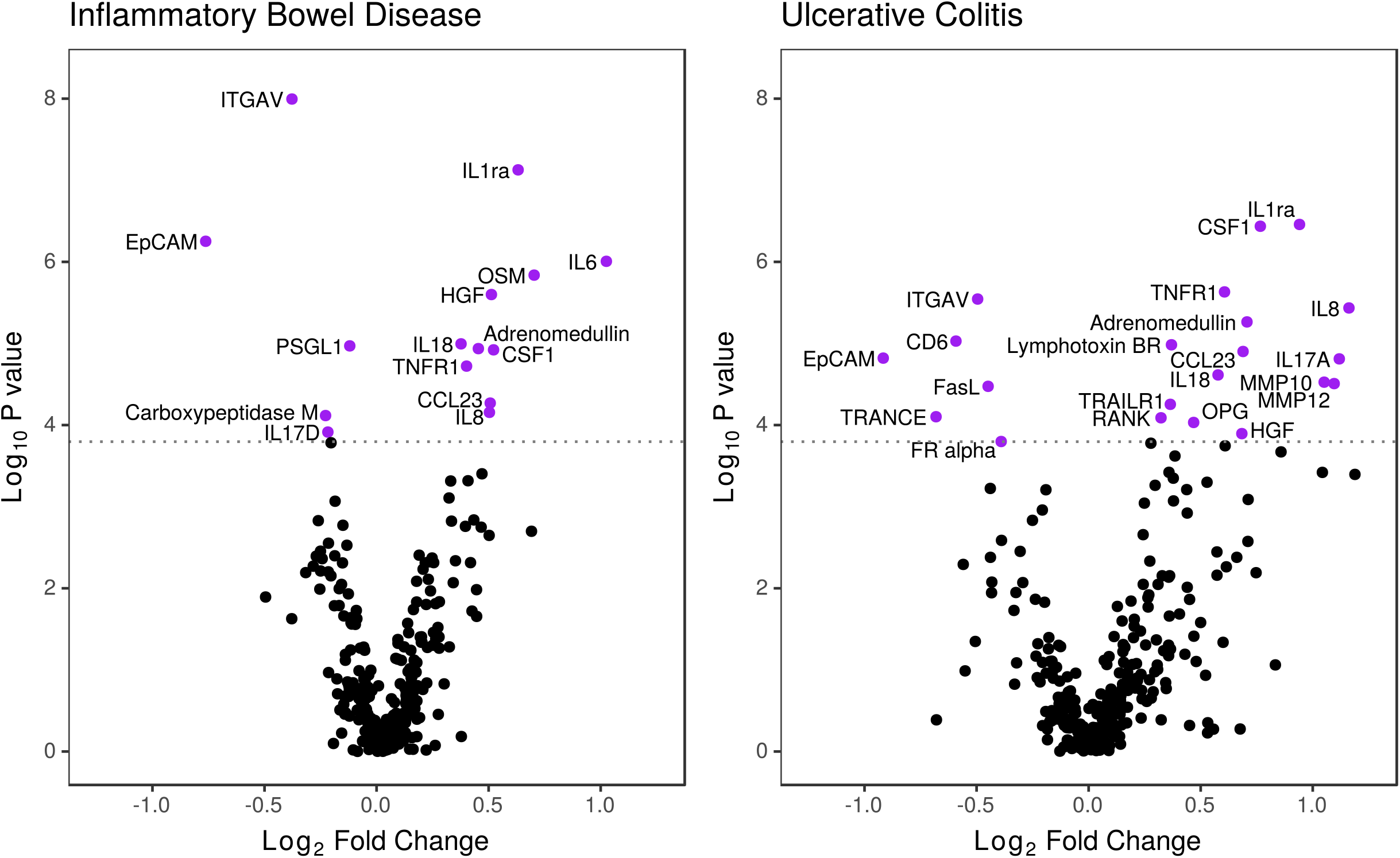
The significance of protein markers in predicting treatment escalation in Inflammatory Bowel Disease and Ulcerative colitis. Significance threshold after Holm correction indicated by dotted line.

**Table 2:**
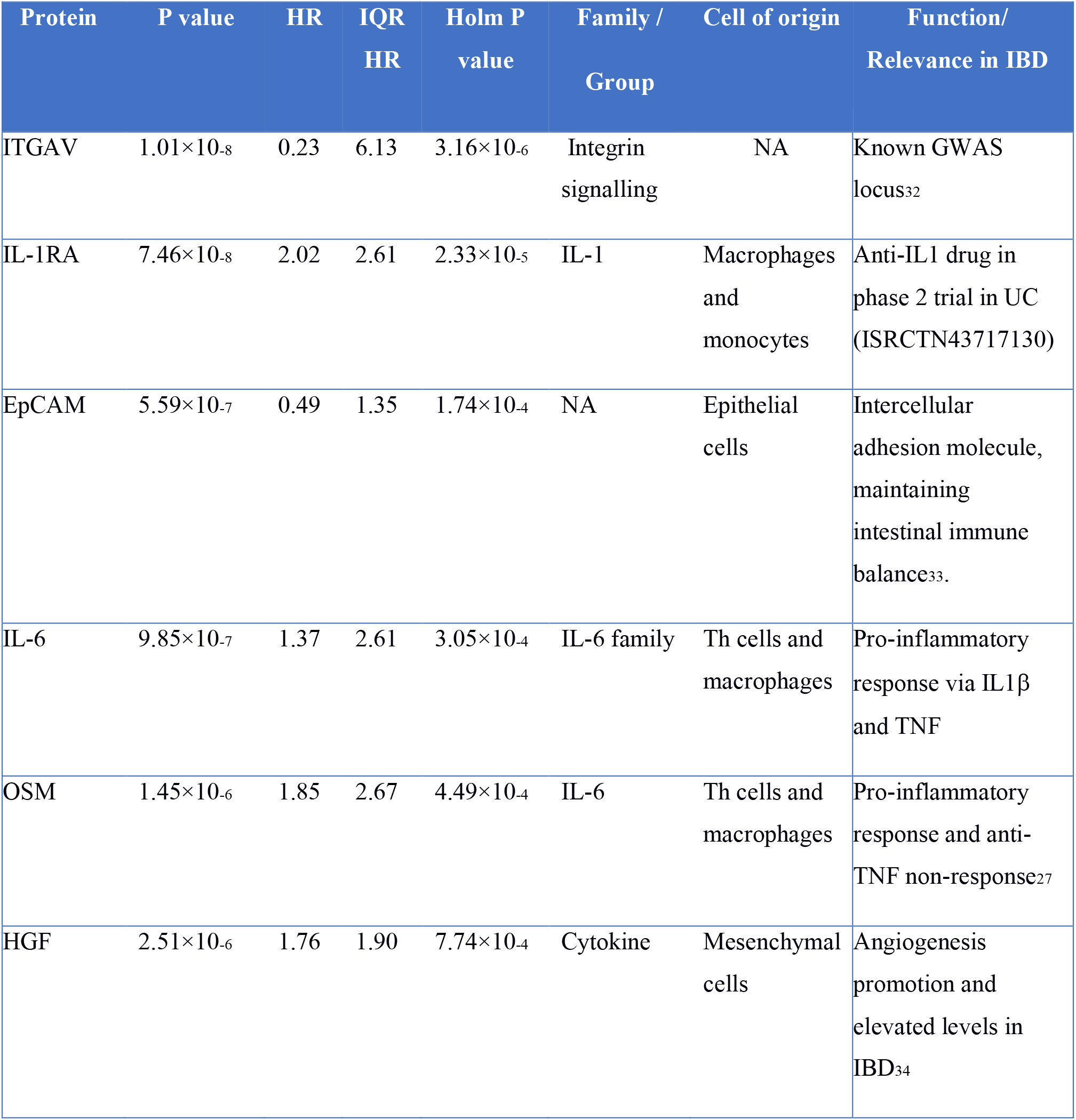
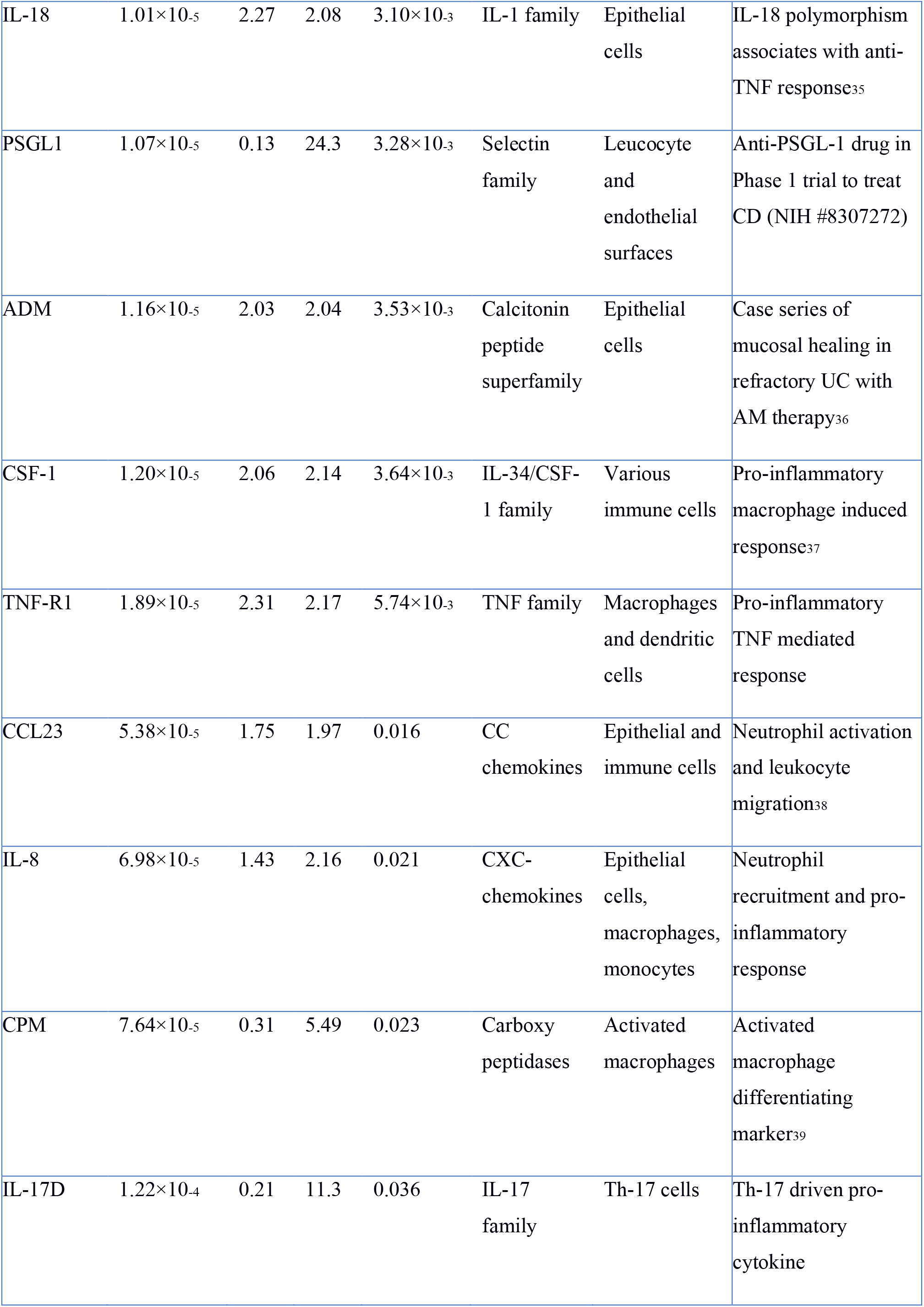
Top 15 proteins associated with escalation in treatment (anti-TNF/ciclosporin and/or surgery) and their associated biology based on the available literature. Holm P represents p values adjusted for multiple testing. HR (hazard ratio) is the relative risk associated with a one unit increase in expression of the relevant protein, IQR HR shows the hazard ratio associated with moving between the 25^th^ and 75^th^ percentile of expression in the direction of increased risk.

### Nested cross-validation stratifies disease sub-groups that associate with treatment escalation

Models to define need for treatment escalation consisting of age, sex, IBD subtype, and PEA-protein expression values were generated in each inner leave-one-out cross-validation loop and tested in the outer loop. The models selected were highly stable. A series of 5-protein models had highest predictive accuracy, with 96% of these models consisting of the same 5 proteins (ITGAV, EpCAM, IL18, SLAMF7, and IL8).

These models defined by cross-validation were 80.0% (CI 75.3-84.7%) accurate (sensitivity 47.6% [CI 35.3-60.0%], specificity 89.6% [85.5-93.7], with a positive likelihood ratio(LR+) 4.59 [2.86-7.36], and negative likelihood ratio(LR-) 0.58 [0.46-0.74]). The high risk group required treatment escalation at 3.9 times the rate of the low risk group (CI 2.4-6.3). FC were higher in patients later requiring treatment escalation (**Table 1**), however this finding was not significant whether analysing CD (p=0.63) and UC (0.09) separately, or in all IBD (p=0.14).

A simple categorisation for all patients as high or low risk may not be the most useful interpretation of the protein expression panels. Subgroups can be identified at particularly high or low risk of aggressive disease tailored to an appropriate level for the intended action to be taken. As an example, identifying the quartiles of patients at highest and lowest risk selects a subset where 52.8% and 5.8% respectively required treatment escalation in the first 18 months of treatment, with a relative risk ratio between groups of 9.1 (**Supplementary Figure S2**).

Although analysing all IBD patients in this cohort together produces models which work in both CD and UC, the accuracy achieved in UC is significantly higher than that in CD (85.1%, CI 79.2-91.0 vs 70.9%, CI 62.4-79.4; χ2 p=0.007). The same analytical approach applied individually to UC and CD produces simpler models (2 and 3 proteins respectively, **Supplementary Figure S3**), with 79.4% (CI 72.8-86.1) accuracy in UC outperforming accuracy in CD(76.4% CI 68.4-84.3). As with the pan-IBD analysis, the probes selected by the inner cross-validation loops were consistent with CD6 and CSF1 in 92% of UC models and LITAF, CPM, and CCL28 in 99, 97, and 88% of CD models respectively **(Table 3).**

**Table 3:**
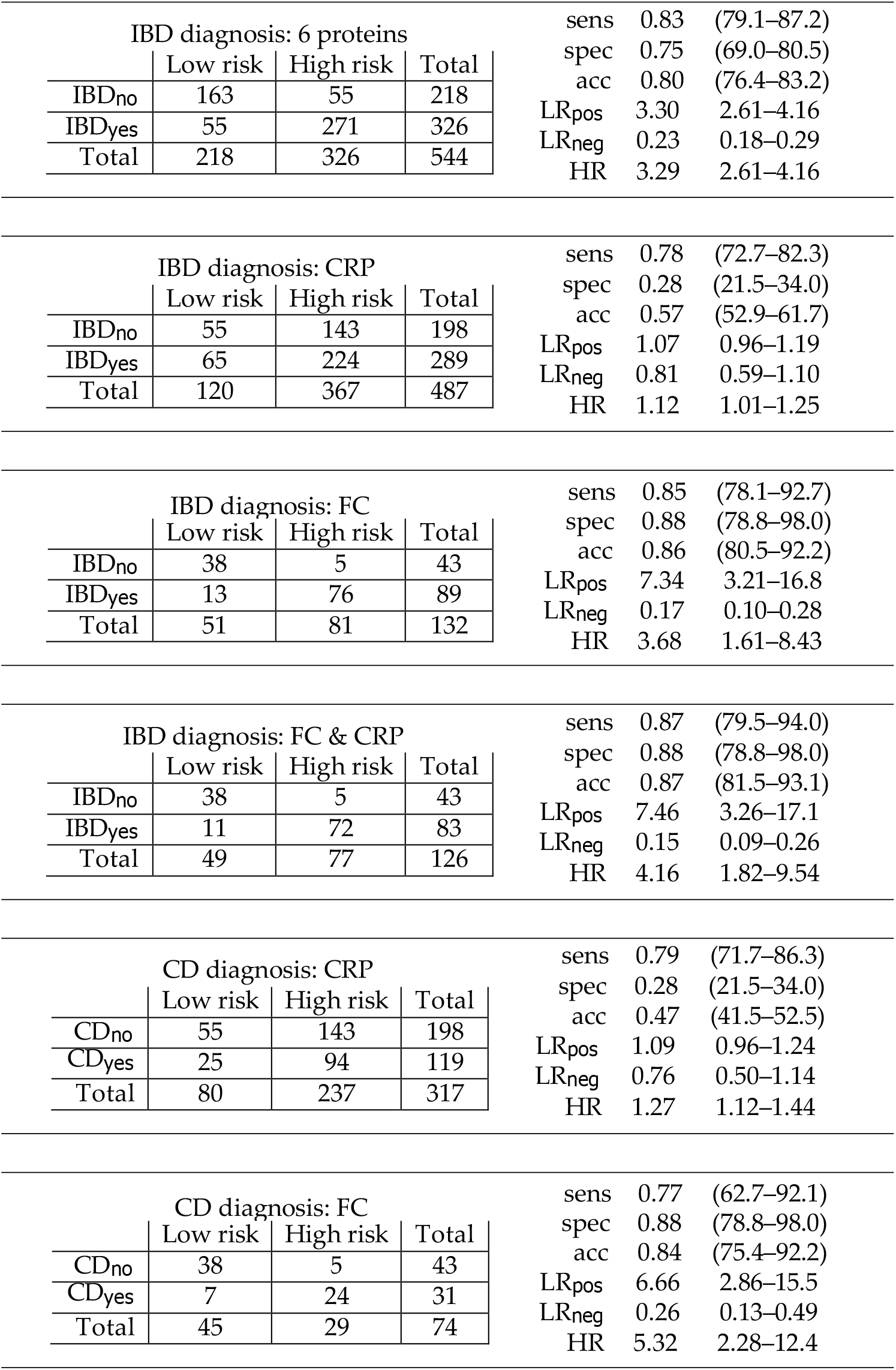
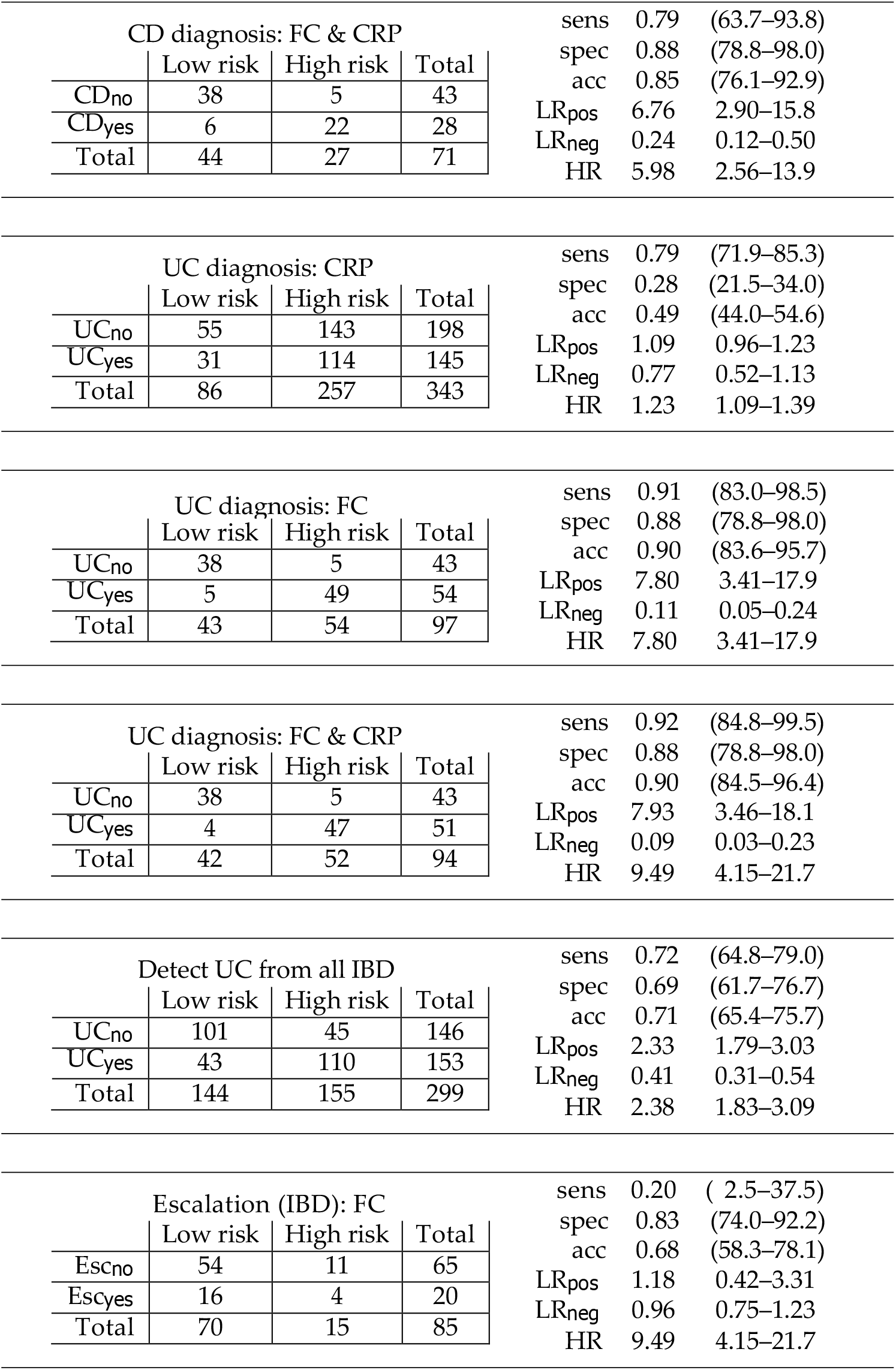
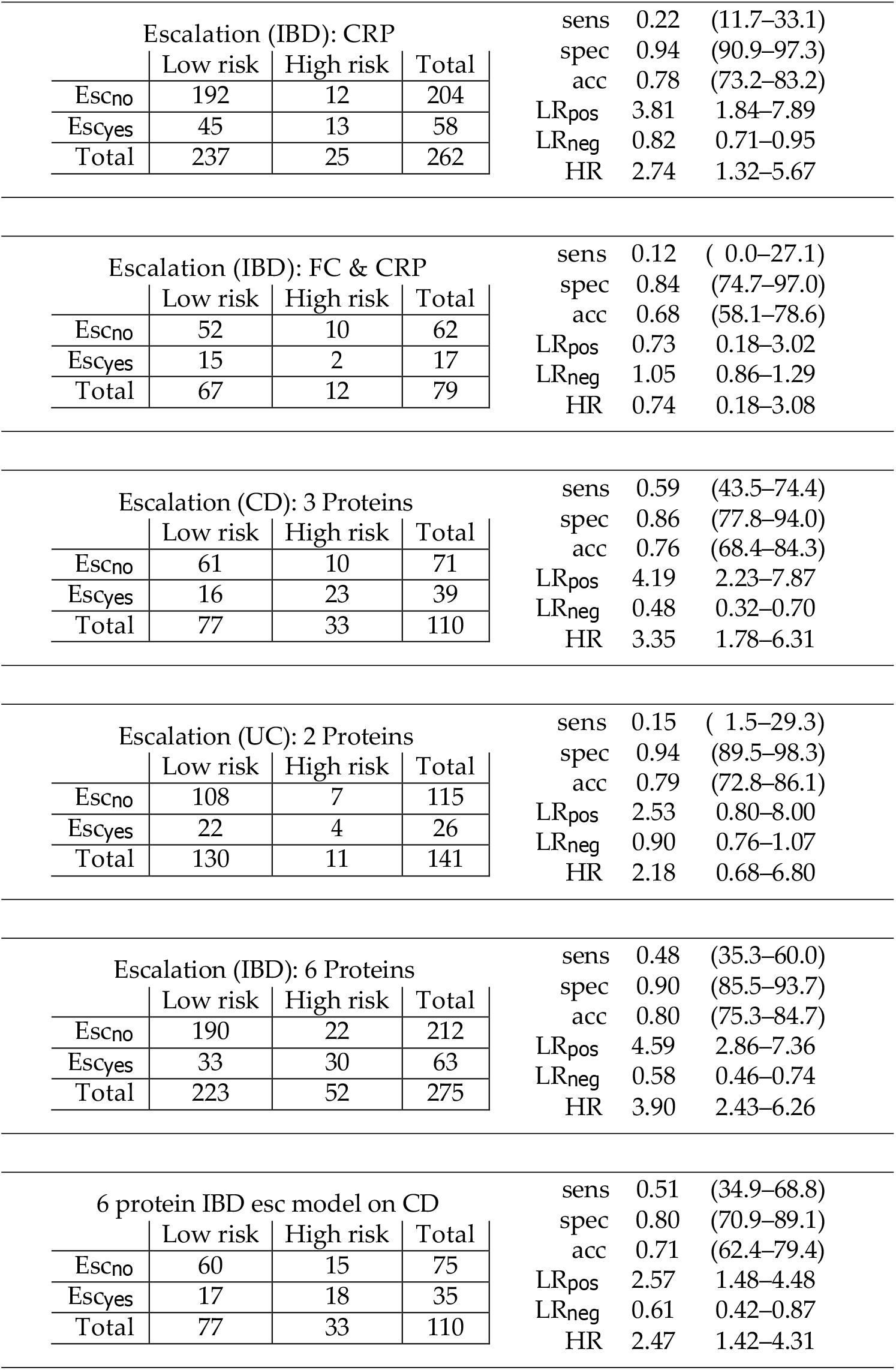
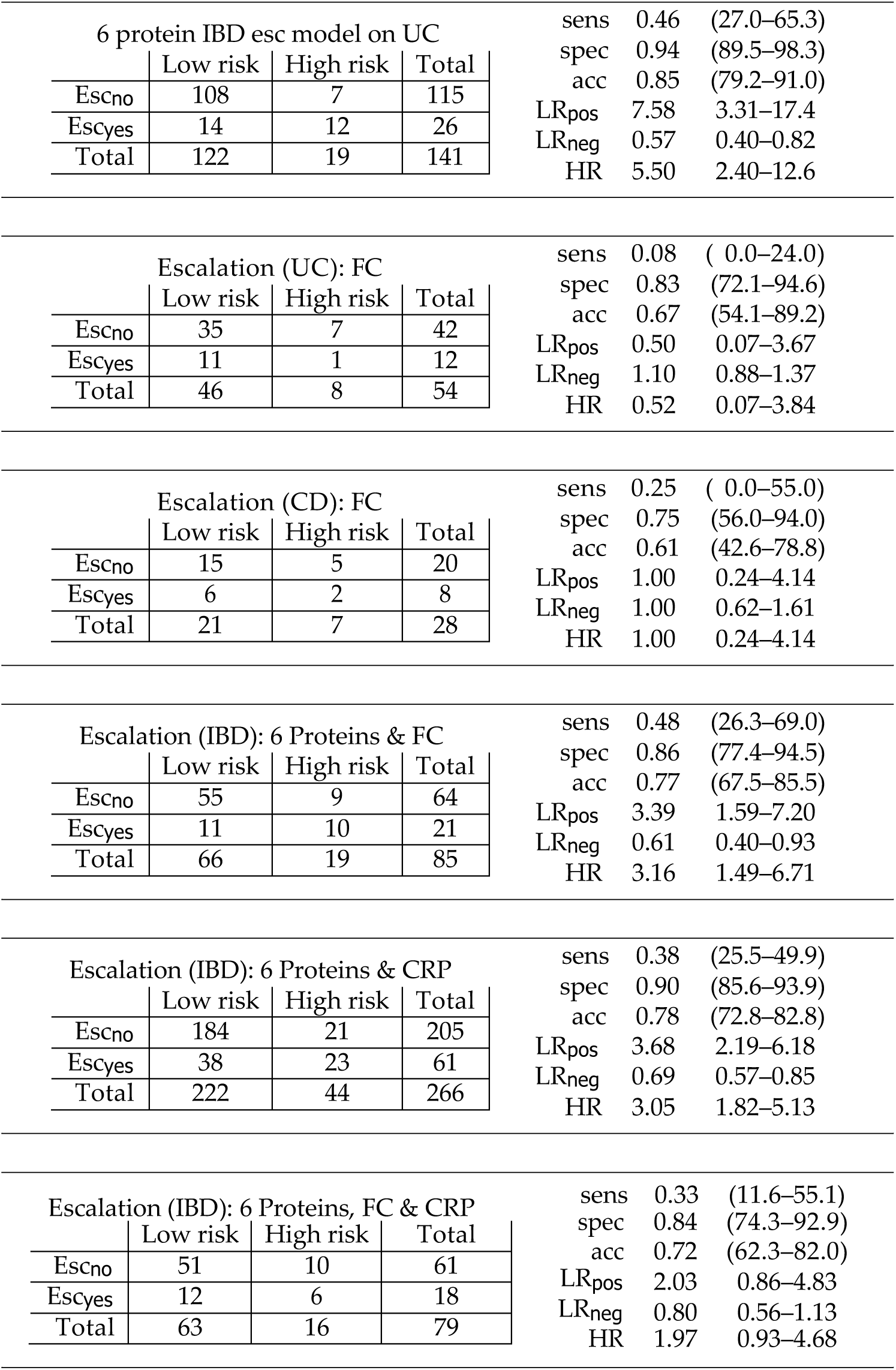
Comparisons of the diagnostic and prognostic performances of Proximity Extension Assay (PEA) based models versus conventional blood tests, faecal markers and clinical predictors of disease. Sens: sensitivity; spec: specificity; acc: accuracy; LRpos: positive likelihood ratio; LRneg: negative likelihood ratio; HR: hazards ratio; IBD: Inflammatory bowel disease; CD: Crohn’s disease; UC: Ulcerative colitis; FC: faecal calprotectin; CRP: high sensitivity c-reactive protein.

| IBD diagnosis: 6 proteins |  |  |  |  |  |  |
| --- | --- | --- | --- | --- | --- | --- |
|  | Low risk | High risk | Total |  |  |  |
| IBD<sub>no</sub> | 163 | 55 | 218 |  |  |  |
| IBD<sub>yes</sub> | 55 | 271 | 326 |  |  |  |
| Total | 218 | 326 | 544 |  |  |  |
|  |  |  |  | sens | 0.83 | (79.1–87.2) |
|  |  |  |  | spec | 0.75 | (69.0–80.5) |
|  |  |  |  | acc | 0.80 | (76.4–83.2) |
|  |  |  |  | LR<sub>pos</sub> | 3.30 | 2.61–4.16 |
|  |  |  |  | LR<sub>neg</sub> | 0.23 | 0.18–0.29 |
|  |  |  |  | HR | 3.29 | 2.61–4.16 |

| IBD diagnosis: CRP |  |  |  |  |  |  |
| --- | --- | --- | --- | --- | --- | --- |
|  | Low risk | High risk | Total |  |  |  |
| IBD<sub>no</sub> | 55 | 143 | 198 |  |  |  |
| IBD<sub>yes</sub> | 65 | 224 | 289 |  |  |  |
| Total | 120 | 367 | 487 |  |  |  |
|  |  |  |  | sens | 0.78 | (72.7–82.3) |
|  |  |  |  | spec | 0.28 | (21.5–34.0) |
|  |  |  |  | acc | 0.57 | (52.9–61.7) |
|  |  |  |  | LR<sub>pos</sub> | 1.07 | 0.96–1.19 |
|  |  |  |  | LR<sub>neg</sub> | 0.81 | 0.59–1.10 |
|  |  |  |  | HR | 1.12 | 1.01–1.25 |

| IBD diagnosis: FC |  |  |  |  |  |  |
| --- | --- | --- | --- | --- | --- | --- |
|  | Low risk | High risk | Total |  |  |  |
| IBD<sub>no</sub> | 38 | 5 | 43 |  |  |  |
| IBD<sub>yes</sub> | 13 | 76 | 89 |  |  |  |
| Total | 51 | 81 | 132 |  |  |  |
|  |  |  |  | sens | 0.85 | (78.1–92.7) |
|  |  |  |  | spec | 0.88 | (78.8–98.0) |
|  |  |  |  | acc | 0.86 | (80.5–92.2) |
|  |  |  |  | LR<sub>pos</sub> | 7.34 | 3.21–16.8 |
|  |  |  |  | LR<sub>neg</sub> | 0.17 | 0.10–0.28 |
|  |  |  |  | HR | 3.68 | 1.61–8.43 |

| IBD diagnosis: FC & CRP |  |  |  |  |  |  |
| --- | --- | --- | --- | --- | --- | --- |
|  | Low risk | High risk | Total |  |  |  |
| IBD<sub>no</sub> | 38 | 5 | 43 |  |  |  |
| IBD<sub>yes</sub> | 11 | 72 | 83 |  |  |  |
| Total | 49 | 77 | 126 |  |  |  |
|  |  |  |  | sens | 0.87 | (79.5–94.0) |
|  |  |  |  | spec | 0.88 | (78.8–98.0) |
|  |  |  |  | acc | 0.87 | (81.5–93.1) |
|  |  |  |  | LR<sub>pos</sub> | 7.46 | 3.26–17.1 |
|  |  |  |  | LR<sub>neg</sub> | 0.15 | 0.09–0.26 |
|  |  |  |  | HR | 4.16 | 1.82–9.54 |

| CD diagnosis: CRP |  |  |  |  |  |  |
| --- | --- | --- | --- | --- | --- | --- |
|  | Low risk | High risk | Total |  |  |  |
| CD<sub>no</sub> | 55 | 143 | 198 |  |  |  |
| CD<sub>yes</sub> | 25 | 94 | 119 |  |  |  |
| Total | 80 | 237 | 317 |  |  |  |
|  |  |  |  | sens | 0.79 | (71.7–86.3) |
|  |  |  |  | spec | 0.28 | (21.5–34.0) |
|  |  |  |  | acc | 0.47 | (41.5–52.5) |
|  |  |  |  | LR<sub>pos</sub> | 1.09 | 0.96–1.24 |
|  |  |  |  | LR<sub>neg</sub> | 0.76 | 0.50–1.14 |
|  |  |  |  | HR | 1.27 | 1.12–1.44 |

| CD diagnosis: FC |  |  |  |  |  |  |
| --- | --- | --- | --- | --- | --- | --- |
|  | Low risk | High risk | Total |  |  |  |
| CD<sub>no</sub> | 38 | 5 | 43 |  |  |  |
| CD<sub>yes</sub> | 7 | 24 | 31 |  |  |  |
| Total | 45 | 29 | 74 |  |  |  |
|  |  |  |  | sens | 0.77 | (62.7–92.1) |
|  |  |  |  | spec | 0.88 | (78.8–98.0) |
|  |  |  |  | acc | 0.84 | (75.4–92.2) |
|  |  |  |  | LR<sub>pos</sub> | 6.66 | 2.86–15.5 |
|  |  |  |  | LR<sub>neg</sub> | 0.26 | 0.13–0.49 |
|  |  |  |  | HR | 5.32 | 2.28–12.4 |

| CD diagnosis: FC & CRP |  |  |  | sens | 0.79 | (63.7–93.8) |
| --- | --- | --- | --- | --- | --- | --- |
|  | Low risk | High risk | Total | spec | 0.88 | (78.8–98.0) |
| CD<sub>no</sub> | 38 | 5 | 43 | acc | 0.85 | (76.1–92.9) |
| CD<sub>yes</sub> | 6 | 22 | 28 | LR<sub>pos</sub> | 6.76 | 2.90–15.8 |
| Total | 44 | 27 | 71 | LR<sub>neg</sub> | 0.24 | 0.12–0.50 |
|  |  |  |  | HR | 5.98 | 2.56–13.9 |

| UC diagnosis: CRP |  |  |  | sens | 0.79 | (71.9–85.3) |
| --- | --- | --- | --- | --- | --- | --- |
|  | Low risk | High risk | Total | spec | 0.28 | (21.5–34.0) |
| UC<sub>no</sub> | 55 | 143 | 198 | acc | 0.49 | (44.0–54.6) |
| UC<sub>yes</sub> | 31 | 114 | 145 | LR<sub>pos</sub> | 1.09 | 0.96–1.23 |
| Total | 86 | 257 | 343 | LR<sub>neg</sub> | 0.77 | 0.52–1.13 |
|  |  |  |  | HR | 1.23 | 1.09–1.39 |

| UC diagnosis: FC |  |  |  | sens | 0.91 | (83.0–98.5) |
| --- | --- | --- | --- | --- | --- | --- |
|  | Low risk | High risk | Total | spec | 0.88 | (78.8–98.0) |
| UC<sub>no</sub> | 38 | 5 | 43 | acc | 0.90 | (83.6–95.7) |
| UC<sub>yes</sub> | 5 | 49 | 54 | LR<sub>pos</sub> | 7.80 | 3.41–17.9 |
| Total | 43 | 54 | 97 | LR<sub>neg</sub> | 0.11 | 0.05–0.24 |
|  |  |  |  | HR | 7.80 | 3.41–17.9 |

| UC diagnosis: FC & CRP |  |  |  | sens | 0.92 | (84.8–99.5) |
| --- | --- | --- | --- | --- | --- | --- |
|  | Low risk | High risk | Total | spec | 0.88 | (78.8–98.0) |
| UC<sub>no</sub> | 38 | 5 | 43 | acc | 0.90 | (84.5–96.4) |
| UC<sub>yes</sub> | 4 | 47 | 51 | LR<sub>pos</sub> | 7.93 | 3.46–18.1 |
| Total | 42 | 52 | 94 | LR<sub>neg</sub> | 0.09 | 0.03–0.23 |
|  |  |  |  | HR | 9.49 | 4.15–21.7 |

| Detect UC from all IBD |  |  |  | sens | 0.72 | (64.8–79.0) |
| --- | --- | --- | --- | --- | --- | --- |
|  | Low risk | High risk | Total | spec | 0.69 | (61.7–76.7) |
| UC<sub>no</sub> | 101 | 45 | 146 | acc | 0.71 | (65.4–75.7) |
| UC<sub>yes</sub> | 43 | 110 | 153 | LR<sub>pos</sub> | 2.33 | 1.79–3.03 |
| Total | 144 | 155 | 299 | LR<sub>neg</sub> | 0.41 | 0.31–0.54 |
|  |  |  |  | HR | 2.38 | 1.83–3.09 |

| Escalation (IBD): FC |  |  |  | sens | 0.20 | ( 2.5–37.5) |
| --- | --- | --- | --- | --- | --- | --- |
|  | Low risk | High risk | Total | spec | 0.83 | (74.0–92.2) |
| Esc<sub>no</sub> | 54 | 11 | 65 | acc | 0.68 | (58.3–78.1) |
| Esc<sub>yes</sub> | 16 | 4 | 20 | LR<sub>pos</sub> | 1.18 | 0.42–3.31 |
| Total | 70 | 15 | 85 | LR<sub>neg</sub> | 0.96 | 0.75–1.23 |
|  |  |  |  | HR | 9.49 | 4.15–21.7 |
| Escalation (IBD): CRP |  |  |  | sens | 0.22 | (11.7–33.1) |
|  | Low risk | High risk | Total | spec | 0.94 | (90.9–97.3) |
| Escno | 192 | 12 | 204 | acc | 0.78 | (73.2–83.2) |
| Escyes | 45 | 13 | 58 | LR<sub>pos</sub> | 3.81 | 1.84–7.89 |
| Total | 237 | 25 | 262 | LR<sub>neg</sub> | 0.82 | 0.71–0.95 |
|  |  |  |  | HR | 2.74 | 1.32–5.67 |
| Escalation (IBD): FC & CRP |  |  |  | sens | 0.12 | ( 0.0–27.1) |
|  | Low risk | High risk | Total | spec | 0.84 | (74.7–97.0) |
| Escno | 52 | 10 | 62 | acc | 0.68 | (58.1–78.6) |
| Escyes | 15 | 2 | 17 | LR<sub>pos</sub> | 0.73 | 0.18–3.02 |
| Total | 67 | 12 | 79 | LR<sub>neg</sub> | 1.05 | 0.86–1.29 |
|  |  |  |  | HR | 0.74 | 0.18–3.08 |
| Escalation (CD): 3 Proteins |  |  |  | sens | 0.59 | (43.5–74.4) |
|  | Low risk | High risk | Total | spec | 0.86 | (77.8–94.0) |
| Escno | 61 | 10 | 71 | acc | 0.76 | (68.4–84.3) |
| Escyes | 16 | 23 | 39 | LR<sub>pos</sub> | 4.19 | 2.23–7.87 |
| Total | 77 | 33 | 110 | LR<sub>neg</sub> | 0.48 | 0.32–0.70 |
|  |  |  |  | HR | 3.35 | 1.78–6.31 |
| Escalation (UC): 2 Proteins |  |  |  | sens | 0.15 | ( 1.5–29.3) |
|  | Low risk | High risk | Total | spec | 0.94 | (89.5–98.3) |
| Escno | 108 | 7 | 115 | acc | 0.79 | (72.8–86.1) |
| Escyes | 22 | 4 | 26 | LR<sub>pos</sub> | 2.53 | 0.80–8.00 |
| Total | 130 | 11 | 141 | LR<sub>neg</sub> | 0.90 | 0.76–1.07 |
|  |  |  |  | HR | 2.18 | 0.68–6.80 |
| Escalation (IBD): 6 Proteins |  |  |  | sens | 0.48 | (35.3–60.0) |
|  | Low risk | High risk | Total | spec | 0.90 | (85.5–93.7) |
| Escno | 190 | 22 | 212 | acc | 0.80 | (75.3–84.7) |
| Escyes | 33 | 30 | 63 | LR<sub>pos</sub> | 4.59 | 2.86–7.36 |
| Total | 223 | 52 | 275 | LR<sub>neg</sub> | 0.58 | 0.46–0.74 |
|  |  |  |  | HR | 3.90 | 2.43–6.26 |
| 6 protein IBD esc model on CD |  |  |  | sens | 0.51 | (34.9–68.8) |
|  | Low risk | High risk | Total | spec | 0.80 | (70.9–89.1) |
| Escno | 60 | 15 | 75 | acc | 0.71 | (62.4–79.4) |
| Escyes | 17 | 18 | 35 | LR<sub>pos</sub> | 2.57 | 1.48–4.48 |
| Total | 77 | 33 | 110 | LR<sub>neg</sub> | 0.61 | 0.42–0.87 |
|  |  |  |  | HR | 2.47 | 1.42–4.31 |
| 6 protein IBD esc model on UC |  |  |  | sens | 0.46 | (27.0–65.3) |
|  | Low risk | High risk | Total | spec | 0.94 | (89.5–98.3) |
| Escno | 108 | 7 | 115 | acc | 0.85 | (79.2–91.0) |
| Escyes | 14 | 12 | 26 | LR<sub>pos</sub> | 7.58 | 3.31–17.4 |
| Total | 122 | 19 | 141 | LR<sub>neg</sub> | 0.57 | 0.40–0.82 |
|  |  |  |  | HR | 5.50 | 2.40–12.6 |
| Escalation (UC): FC |  |  |  | sens | 0.08 | ( 0.0–24.0) |
|  | Low risk | High risk | Total | spec | 0.83 | (72.1–94.6) |
| Escno | 35 | 7 | 42 | acc | 0.67 | (54.1–89.2) |
| Escyes | 11 | 1 | 12 | LR<sub>pos</sub> | 0.50 | 0.07–3.67 |
| Total | 46 | 8 | 54 | LR<sub>neg</sub> | 1.10 | 0.88–1.37 |
|  |  |  |  | HR | 0.52 | 0.07–3.84 |
| Escalation (CD): FC |  |  |  | sens | 0.25 | ( 0.0–55.0) |
|  | Low risk | High risk | Total | spec | 0.75 | (56.0–94.0) |
| Escno | 15 | 5 | 20 | acc | 0.61 | (42.6–78.8) |
| Escyes | 6 | 2 | 8 | LR<sub>pos</sub> | 1.00 | 0.24–4.14 |
| Total | 21 | 7 | 28 | LR<sub>neg</sub> | 1.00 | 0.62–1.61 |
|  |  |  |  | HR | 1.00 | 0.24–4.14 |
| Escalation (IBD): 6 Proteins & FC |  |  |  | sens | 0.48 | (26.3–69.0) |
|  | Low risk | High risk | Total | spec | 0.86 | (77.4–94.5) |
| Escno | 55 | 9 | 64 | acc | 0.77 | (67.5–85.5) |
| Escyes | 11 | 10 | 21 | LR<sub>pos</sub> | 3.39 | 1.59–7.20 |
| Total | 66 | 19 | 85 | LR<sub>neg</sub> | 0.61 | 0.40–0.93 |
|  |  |  |  | HR | 3.16 | 1.49–6.71 |
| Escalation (IBD): 6 Proteins & CRP |  |  |  | sens | 0.38 | (25.5–49.9) |
|  | Low risk | High risk | Total | spec | 0.90 | (85.6–93.9) |
| Escno | 184 | 21 | 205 | acc | 0.78 | (72.8–82.8) |
| Escyes | 38 | 23 | 61 | LR<sub>pos</sub> | 3.68 | 2.19–6.18 |
| Total | 222 | 44 | 266 | LR<sub>neg</sub> | 0.69 | 0.57–0.85 |
|  |  |  |  | HR | 3.05 | 1.82–5.13 |
| Escalation (IBD): 6 Proteins, FC & CRP |  |  |  | sens | 0.33 | (11.6–55.1) |
|  | Low risk | High risk | Total | spec | 0.84 | (74.3–92.9) |
| Escno | 51 | 10 | 61 | acc | 0.72 | (62.3–82.0) |
| Escyes | 12 | 6 | 18 | LR<sub>pos</sub> | 2.03 | 0.86–4.83 |
| Total | 63 | 16 | 79 | LR<sub>neg</sub> | 0.80 | 0.56–1.13 |
|  |  |  |  | HR | 1.97 | 0.93–4.68 |

### Performance of PEA prognostic models against conventional predictors of escalation

We compared the performance of PEA based prognostic proteins to currently available blood and faecal biomarkers and clinical predictors in IBD and its subtypes; these are summarised in Table 3. A Cox model trained with FC was highly specific but performed poorly at positively identifying patients who required treatment escalation (sensitivity 20.0%, CI 2.5-37.5; 8.3%, CI 0.0-24.0 in UC and 25%, CI 0.0-55.0 in CD) and suffered from poor uptake with only 85 FC results available for analysis. The performance of the PEA model is comparable to hsCRP (HR 2.74, CI:1.32-5.67 vs 6-protein model HR 3.90, CI:2.43-6.26). It is worth noting however that 149 patients had an hsCRP within the normal range(<5mg/mL).

A combined FC and hsCRP model has a poor performance at predicting escalation (HR 0.74, CI:0.18-3.08). Clinical predictors such as non-B1 behaviour or perianal disease in CD, and SCCAI or HBI scores did not significantly associate with treatment escalation, though pancolitis in UC did (uncorrected p=0.002).

Compared to the overall PEA-protein model accuracy of 80.0%, the addition of FC, CRP, or both did not improve model performance yielding accuracies of 76.5% (CI 67.5-85.5) 77.8% (72.8-82.8), and 72.2% (62.3-82.0) respectively, neither did the addition of any phenotypic characteristic such as pancolitis in UC or perianal disease in CD. We also performed correlation analyses of the top protein markers with proteins associated with IBD, hsCRP, albumin, and FC and these are summarised in Supplementary Figure S4.

### Circulating proteins associate with germline variation

It has been shown that expression of proteins associate with germline variation, mainly in the cis regions of their encoding genes^23^. We explored the influence of germline variation on the expression of key IBD diagnostic and prognostic proteins identified in our analysis. We used linear regression models with age and sex as covariates, to analyse SNPs (MAF >0.1) correlated with protein expression, revealing 769 significant cis pQTLs (Holm corrected) affecting 51 proteins. These included 59 significant cis pQTLs affecting 9 proteins with significant expression changes associated with IBD, (**Supplementary Figure S5, Supplementary Table 9**), and 35 pQTLs affecting proteins implicated in disease course (**Supplementary Figures S6**). Vascular Endothelial Growth Factor-A (VEGF-A) showed the most significant association with genotype (lead SNP rs7767396; effect (β) −0.42; MAF=0.46; p=8.7×10^−18^) with a total of 6 significant SNP associations and 14 SNPs in linkage disequilibrium with rs7767396.

Among the proteins individually significantly associated with aggressive disease (**Table 2**) or frequently selected in the multi-protein models for aggressive disease significant pQTLs were found in CD6, RANK and SLAMF7 (**Supplementary Figure S6**), in addition to the findings described in CCL23 above (**Supplementary Figure S5**).

## Discussion

With advances in clinical care in IBD, it is widely recognised that there is a need for biomarkers that provide accurate diagnostic and prognostic testing in IBD. The key innovation in this study is the design and evaluation of a novel multi-protein panels in newly diagnosed IBD, chosen a priori on the basis of known or suspected involvement in pathogenesis. The results substantiate the involvement of key pathways in pathogenesis, as well as provide targets for therapy. Importantly, we demonstrate that this strategy of biomarker discovery is feasible in diagnosis and in predicting treatment escalation in CD and UC.

A panel of 6 proteins had 79.8 % accuracy, 83.1% sensitivity, and 74.8% specificity at differentiating IBD from controls. Whilst FC did outperform this panel (86.4% accuracy, 85.4% sensitivity, 88.4% specificity), uptake was low, overall with patient acceptability a major limiting factor. We suggest a serum protein biomarker panel could prove clinically useful given this widely recognised limitations of FC testing in clinic^10,11^.

Of the 66 differentially expressed proteins in IBD, 9 demonstrated germline variation, VEGF-A being the most significant pQTL. Weaker correlations between protein expression and genetic variation were observed in 4 of the proteins that predicted treatment escalation including CCL23, RANK, CD6 and SLAM7. It is yet to be determined whether these genetic associations are causal in both disease onset and course and our study provide a resource to investigate these associations further.

The greatest unmet need is for biomarkers that can determine disease activity, behaviour and extent, and most critically to predict response to treatment. In our dataset, we have been able to characterise and rigorously cross-validate models involving a limited number of proteins that predict disease course. The role of biomarkers in predicting the disease course has been the focus of many studies^2–7,24,25^, including our own parallel studies of glycomic and methylation profiling in the EC-funded consortia^24,26^. Lee *et al* identified expression profiles of T cell exhaustion in CD8 T cells that predicted treatment escalation in IBD^3^, defining escalation as the need for 2 or more immunosuppressants and/or surgery after initial disease remission. A multi-gene signature predicting need for escalation using these original criteria has been proposed by this team in UC (HR 3.1, 95% CI: 1.25-7.72, p=0.02) and CD (HR 2.7; CI: 1.32-5.34, p=0.01)^7^. This signature differs from the original profile of T cell exhaustion. Other studies focus on mucosal healing, response to biological agents, and development of fistulising or stricturing complications as end-points – all valid in context.

In this study we decided to use more stringent criteria for escalation than those used in defining the transcriptional profile. We highlight need for biologics or ciclosporin or surgical resection, rather than introduction of immunosuppression per se. This decision regarding end-point relates principally to the variable threshold for initiating immuno-modulators, which in practice have often been used as first-line therapy in CD. Our oligo-protein panels have the potential for clinical translation with significant practical benefits including the simplicity of the assay, and the ability to multiplex proteins using only 1μL of serum.

It is noteworthy that the key prognostic proteins identified relate to pathways independent of TNF signalling. OSM is a pro-inflammatory cytokine that promotes production of IL-6 to attract immune cells to the site of inflammation^27^ and its intestinal expression in IBD has been shown to predict anti-TNF non-response in IBD^27^. We report that circulating levels of both IL-6 and OSM can predict treatment escalation in IBD. Similarly, we demonstrate the involvement of other pathways that predict disease course (**Table 2, Supplementary Figure S7**). Of particular relevance are the proteins that show poor correlation with conventional inflammatory markers including hsCRP (**Supplementary Figure 4**), in particular PSGL-1. This protein is a P-selectin glycoprotein ligand that is expressed on the surface of most immune cells and facilitates immune cell trafficking across the endothelium^28,29^. Drug targeting PSGL-1 is currently in phase 1 trial for the treatment of CD (NIH #8307272). Future studies examining the performance of these markers in predicting response to therapy are now needed.

We recognise that clinical decisions and timing on treatment escalations may vary across centres. In this study all sites utilised a ‘step-up approach’ to treatment escalation, rather than a top-down approach. In this respect the clinical management is similar across centres and the consistency of our biomarker profile in predicting need for escalation across centres is especially noteworthy. Our study was not designed to detect the association between prognosis and endoscopic activity. Recently, a protein based endoscopic healing index (EHI) has been reported that incorporates 13 proteins and performs at par with FC in predicting endoscopic disease remission (validation cohort AUROC, 0.803 for EHI vs AUROC, 0.854 for FC; P = .298); highlighting the translational potential of blood-based protein biomarkers in IBD^30^. Our prognostic protein model performs at par with conventional blood tests such as hsCRP. Clinicians often make treatment decisions based on these biomarkers, confounding the performance of hsCRP and albumin in prognostication. It is however worth noting that 149 patients with IBD had an hsCRP within the normal range (<5mg/mL). Furthermore, our protein markers still remain significant predictors of treatment escalation, independent of clinical confounders. We have utilised nested leave-one-out cross-validation which is acknowledged to produce an unbiased estimate of true error when properly nested so that the entire feature selection and parameter tuning process takes place without reference to the left out samples^31^. Further validation is now needed to replicate our findings in other large multi-centre inception studies. The significance and impact of our analysis are strengthened by the pre-established evidence for these proteins in IBD or IBD-related pathways. This is however the largest inception cohort recruited in biomarker studies in adult IBD to date, allowing robust modelling and rigorous application.

With advances in IBD therapeutics, future challenges will include tailoring therapies based on individual disease biology. Our data provide an insight into the importance of molecular characterisation of patients with IBD at diagnosis to tailor medical therapies. With the setup and initiation of the biomarker-stratified PROFILE trial in CD 25, the aspiration is that stratification with multi-omic biomarkers based on underlying disease mechanisms may enable personalised therapeutics.

## Author Contributions

Study design RK, JH, MDA, MV, JS. Patient recruitment and sample processing NTV, RK, NAK, DB, SV, ATA. Experimental work OA, FH, CP, RK, NTV, ATA, NAK. Data Analysis RK, NAK, ATA, DR, JL, DB. RK and ATA wrote the manuscript. All authors were involved in critical review, editing, revision and approval of the final manuscript.

---

**Supplementary Figure S1:** Heatmap showing correlation coefficients of PEA assays with high sensitivity C-reactive protein (hsCRP), albumin (Alb) in the entire cohort. Colour shows absolute correlation and figures show relative correlation.

**Supplementary Figure S2:** Each subsection represents the results from labelling a proportion of the population as low (x axis) and high risk (y axis). Within each subsection the top left and bottom right numbers denote the percentage of the identified group requiring escalated treatment in the high and low risk groups respectively. The top right number in each subsection represents the relative risk between groups. The equivalent results obtained by categorisation based on optimum ROC thresholds are: 22.9% high risk with 47.6% escalation, 77.1% low risk with 10.4% escalation, relative risk = 4.6 (95% CI 2.9-7.4)

**Supplementary Figure S3:** Kaplan-Meier survival graphs showing stratification of CD and UC patients by 3 and 2 PEA assays respectively into groups at high and low risk of treatment escalation.

**Supplementary Figure S4:** Heatmap showing hierarchical clustering by absolute correlation (Spearman) between top 66 differentially expressed proteins, escalation-associated proteins, high sensitivity C-reactive protein (hsCRP), albumin(Alb), and faecal calprotectin (FCP). Colour shows absolute correlation and figures show relative correlation.

**Supplementary Figure S5:** All proteins where there is a significant association between expression and disease status, and cis pQTLs with SNPs within 300Kb.

**Supplementary Figure S6:** All proteins associated with escalation (individually or in the stepwise constructed models) with cis pQTLs with SNPs within 300Kb.

**Supplementary Figure S7**: Heatmap summarising the associations of the top differentially expressed prognostic proteins to the cell-specific Cap Analysis of Gene Expression (CAGE) peaks derived using the FANTOM 5 dataset^40^. Markers that predict disease course, cluster into distinct groups based on their expression in cell lines within the FANTOM-5 dataset. These include protein markers that associate with the innate immune system such as macrophages and mast cells (IL-18, IL-1RA, CCL23, CSF-1) and a distinct group of proteins that are primarily expressed in monocytes (IL-6, ITGAV).

## References

1. Boyapati RK., Kalla R., Satsangi J., Ho G-T. Biomarkers in Search of Precision Medicine in IBD. Am J Gastroenterol 2016;111(12):1682–90. Doi: 10.1038/ajg.2016.441.

2. Kalla R., Kennedy NA., Ventham NT., Boyapati RK., Adams AT., Nimmo ER., et al. Serum Calprotectin: A Novel Diagnostic and Prognostic Marker in Inflammatory Bowel Diseases. Am J Gastroenterol 2016;111(12):1796–805. Doi: 10.1038/ajg.2016.342.

3. Lee JC., Lyons P a., McKinney EF., Sowerby JM., Carr EJ., Bredin F., et al. Gene expression profiling of CD8+ T cells predicts prognosis in patients with Crohn disease and ulcerative colitis. J Clin Invest 2011;121(10):4170–9. Doi: 10.1172/JCI59255.

4. Lee JC., Biasci D., Roberts R., Gearry RB., Mansfield JC., Ahmad T., et al. Genome-wide association study identifies distinct genetic contributions to prognosis and susceptibility in Crohn’s disease. Nat Genet 2017;49(2):262–8. Doi: 10.1038/ng.3755.

5. Kugathasan S., Denson LA., Walters TD., Kim M-O., Marigorta UM., Schirmer M., et al. Prediction of complicated disease course for children newly diagnosed with Crohn’s disease: a multicentre inception cohort study. Lancet (London, England) 2017;389(10080):1710–8. Doi: 10.1016/S0140-6736(17)30317-3.

6. Marigorta UM., Denson LA., Hyams JS., Mondal K., Prince J., Walters TD., et al. Transcriptional risk scores link GWAS to eQTLs and predict complications in Crohn’s disease. Nat Genet 2017;49(10):1517–21. Doi: 10.1038/ng.3936.

7. Biasci D., Lee JC., Noor NM., Pombal DR., Hou M., Lewis N., et al. A blood-based prognostic biomarker in IBD. Gut 2019;68(8):1386–95. Doi: 10.1136/gutjnl-2019-318343.

8. Colombel J-F., Panaccione R., Bossuyt P., Lukas M., Baert F., Vaňásek T., et al. Effect of tight control management on Crohn’s disease (CALM): a multicentre, randomised, controlled phase 3 trial. Lancet 2017;390(10114):2779–89. Doi: 10.1016/S0140-6736(17)32641-7.

9. van Rheenen PF., Van de Vijver E., Fidler V. Faecal calprotectin for screening of patients with suspected inflammatory bowel disease: diagnostic meta-analysis. BMJ 2010;341:c3369.

10. Kalla R., Boyapati R., Vatn S., Hijos G., Crooks B., Moore GT., et al. Patients’ perceptions of faecal calprotectin testing in inflammatory bowel disease: results from a prospective multicentre patient-based survey. Scand J Gastroenterol 2018:1–6. Doi: 10.1080/00365521.2018.1527394.

11. Maréchal C., Aimone-Gastin I., Baumann C., Dirrenberger B., Guéant J-L., Peyrin-Biroulet L. Compliance with the faecal calprotectin test in patients with inflammatory bowel disease. United Eur Gastroenterol J 2017:205064061668651. Doi: 10.1177/2050640616686517.

12. Assarsson E., Lundberg M., Holmquist G., Björkesten J., Thorsen SB., Ekman D., et al. Homogenous 96-plex PEA immunoassay exhibiting high sensitivity, specificity, and excellent scalability. PLoS One 2014;9(4):e95192. Doi: 10.1371/journal.pone.0095192.

13. Lundberg M., Eriksson A., Tran B., Assarsson E., Fredriksson S. Homogeneous antibody-based proximity extension assays provide sensitive and specific detection of low-abundant proteins in human blood. Nucleic Acids Res 2011;39(15):e102. Doi: 10.1093/nar/gkr424.

14. Lennard-Jones JE. Classification of inflammatory bowel disease. Scand J Gastroenterol Suppl 1989;170:2–6; discussion 16-9.

15. Satsangi J., Silverberg MS., Vermeire S., Colombel J-F. The Montreal classification of inflammatory bowel disease: controversies, consensus, and implications. Gut 2006;55(6):749–53. Doi: 10.1136/gut.2005.082909.

16. Levine A., Griffiths A., Markowitz J., Wilson DC., Turner D., Russell RK., et al. Pediatric modification of the Montreal classification for inflammatory bowel disease: the Paris classification. Inflamm Bowel Dis 2011;17(6): 1314–21. Doi: 10.1002/ibd.21493.

17. Jostins L., Ripke S., Weersma RK., Duerr RH., McGovern DP., Hui KY., et al. Host-microbe interactions have shaped the genetic architecture of inflammatory bowel disease. Nature 2012;491(7422):119–24. Doi: 10.1038/nature11582.

18. Franke A., Mcgovern DPB., Barrett JC., Wang K., Graham L., Ahmad T., et al. Meta-Analysis Increases to 71 the Tally of Confirmed Crohn’s Disease Susceptibility Loci. Nat Genet 2010;42(12):1118–25. Doi: 10.1038/ng.717.Meta-Analysis.

19. Lind L., Ärnlöv J., Lindahl B., Siegbahn A., Sundström J., Ingelsson E. Use of a proximity extension assay proteomics chip to discover new biomarkers for human atherosclerosis. Atherosclerosis 2015;242(1):205–10. Doi: 10.1016/j.atherosclerosis.2015.07.023.

20. Bezanson J., Edelman A., Karpinski S., Shah VB. Julia: A Fresh Approach to Numerical Computing. SIAM Rev 2017;59(1):65–98. Doi: 10.1137/141000671.

21. Holm S. A simple sequentially rejective multiple test procedure. Scand J Stat 1979;6(2):65–70.

22. Shabalin AA. Matrix eQTL: ultra fast eQTL analysis via large matrix operations. Bioinformatics 2012;28(10):1353–8. Doi: 10.1093/bioinformatics/bts163.

23. Sun BB., Maranville JC., Peters JE., Stacey D., Staley JR., Blackshaw J., et al. Genomic atlas of the human plasma proteome. Nature 2018;558(7708):73–9. Doi: 10.1038/s41586-018-0175-2.

24. Clerc F., Novokmet M., Dotz V., Reiding KR., de Haan N., Kammeijer GSM., et al. Plasma N-Glycan Signatures Are Associated With Features of Inflammatory Bowel Diseases. Gastroenterology 2018;155(3):829–43. Doi: 10.1053/j.gastro.2018.05.030.

25. Parkes M., Noor NM., Dowling F., Leung H., Bond S., Whitehead L., et al. PRedicting Outcomes For Crohn’s dIsease using a moLecular biomarkEr (PROFILE): protocol for a multicentre, randomised, biomarker-stratified trial. BMJ Open 2018;8(12):e026767. Doi: 10.1136/bmjopen-2018-026767.

26. Ventham NT., Kennedy NA., Adams AT., Kalla R., Heath S., O’Leary KR., et al. Integrative epigenome-wide analysis demonstrates that DNA methylation may mediate genetic risk in inflammatory bowel disease. Nat Commun 2016;7:13507. Doi: 10.1038/ncomms13507.

27. West NR., Hegazy AN., Owens BMJ., Bullers SJ., Linggi B., Buonocore S., et al. Oncostatin M drives intestinal inflammation and predicts response to tumor necrosis factor-neutralizing therapy in patients with inflammatory bowel disease. Nat Med 2017;23(5):579–89. Doi: 10.1038/nm.4307.

28. Guyer DA., Moore KL., Lynam EB., Schammel CM., Rogelj S., McEver RP., et al. P-selectin glycoprotein ligand-1 (PSGL-1) is a ligand for L-selectin in neutrophil aggregation. Blood 1996;88(7):2415–21.

29. Brown JB., Cheresh P., Zhang Z., Ryu H., Managlia E., Barrett TA. P-selectin glycoprotein ligand-1 is needed for sequential recruitment of T-helper 1 (Th1) and local generation of Th17 T cells in dextran sodium sulfate (DSS) colitis. Inflamm Bowel Dis 2012;18(2):323–32. Doi: 10.1002/ibd.21779.

30. D’Haens G., Kelly O., Battat R., Silverberg MS., Laharie D., Louis E., et al. Development and Validation of a Test to Monitor Endoscopic Activity in Patients With Crohn’s Disease Based on Serum Levels of Proteins. Gastroenterology 2020;158(3):515–526.e10. Doi: 10.1053/j.gastro.2019.10.034.

31. Lachenbruch PA., Mickey MR. Estimation of Error Rates in Discriminant Analysis. Technometrics 1968;10(1):1–11. Doi: 10.1080/00401706.1968.10490530.

32. de Lange KM., Moutsianas L., Lee JC., Lamb CA., Luo Y., Kennedy NA., et al. Genome-wide association study implicates immune activation of multiple integrin genes in inflammatory bowel disease. Nat Genet 2017;49(2):256–61. Doi: 10.1038/ng.3760.

33. Jiang L., Shen Y., Guo D., Yang D., Liu J., Fei X., et al. EpCAM-dependent extracellular vesicles from intestinal epithelial cells maintain intestinal tract immune balance. Nat Commun 2016;7:13045. Doi: 10.1038/ncomms13045.

34. Srivastava M., Zurakowski D., Cheifetz P., Leichtner A., Bousvaros A. Elevated serum hepatocyte growth factor in children and young adults with inflammatory bowel disease. J Pediatr Gastroenterol Nutr 2001;33(5):548–53.

35. Bank S., Julsgaard M., Abed OK., Burisch J., Broder Brodersen J., Pedersen NK., et al. Polymorphisms in the NFkB, TNF-alpha, IL-1beta, and IL-18 pathways are associated with response to anti-TNF therapy in Danish patients with inflammatory bowel disease. Aliment Pharmacol Ther 2019;49(7):890–903. Doi: 10.1111/apt.15187.

36. Ashizuka S., Inatsu H., Kita T., Kitamura K. Adrenomedullin Therapy in Patients with Refractory Ulcerative Colitis: A Case Series. Dig Dis Sci 2016;61(3):872–80. Doi: 10.1007/s10620-015-3917-0.

37. Marshall D., Cameron J., Lightwood D., Lawson ADG. Blockade of colony stimulating factor-1 (CSF-I) leads to inhibition of DSS-induced colitis. Inflamm Bowel Dis 2007;13(2):219–24. Doi: 10.1002/ibd.20055.

38. Singh UP., Singh NP., Murphy EA., Price RL., Fayad R., Nagarkatti M., et al. Chemokine and cytokine levels in inflammatory bowel disease patients. Cytokine 2016;77:44–9. Doi: 10.1016/j.cyto.2015.10.008.

39. Tsakiris I., Torocsik D., Gyongyosi A., Dozsa A., Szatmari I., Szanto A., et al. Carboxypeptidase-M is regulated by lipids and CSFs in macrophages and dendritic cells and expressed selectively in tissue granulomas and foam cells. Lab Invest 2012;92(3):345–61. Doi: 10.1038/labinvest.2011.168.

40. FANTOM Consortium and the RIKEN PMI and CLST (DGT)., Forrest ARR., Kawaji H., Rehli M., Baillie JK., de Hoon MJL., et al. A promoter-level mammalian expression atlas. Nature 2014;507(7493):462–70. Doi: 10.1038/nature13182.

