## Supplementary figures and images for "Serum proteomic profiling at diagnosis predicts clinical course, and need for intensification of treatment in inflammatory bowel disease"

### Fig S1

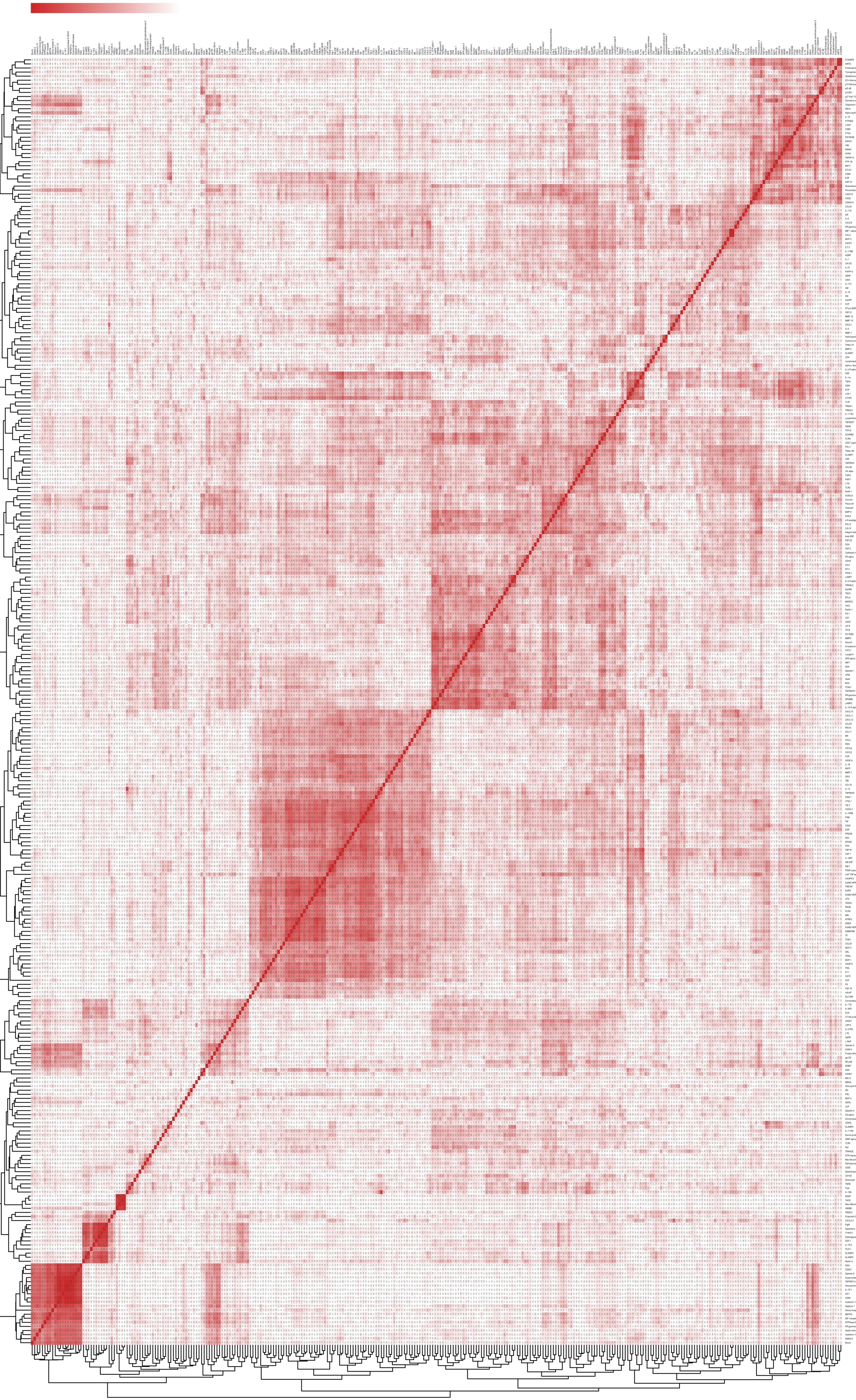

### Fig S2

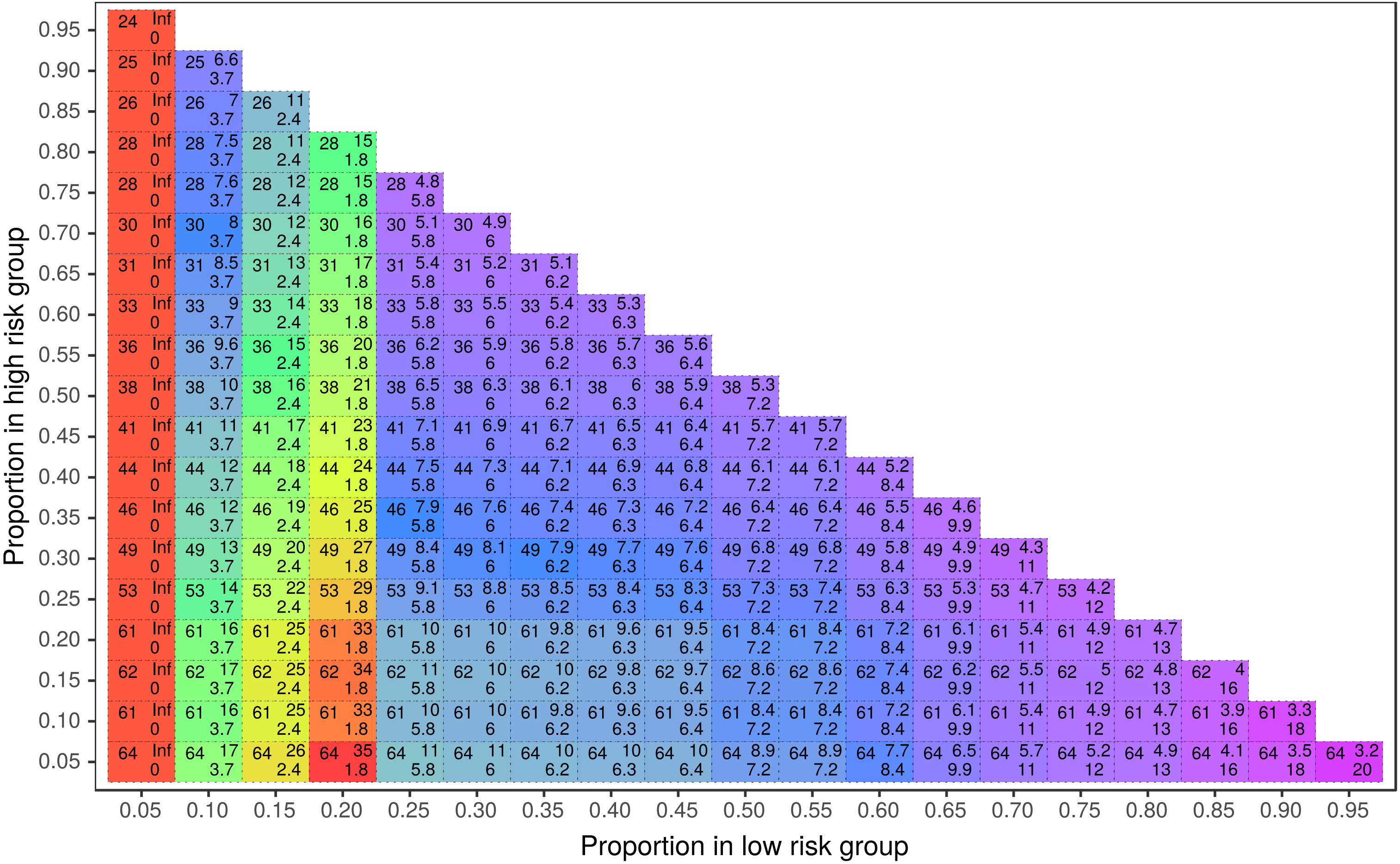

### Fig S3

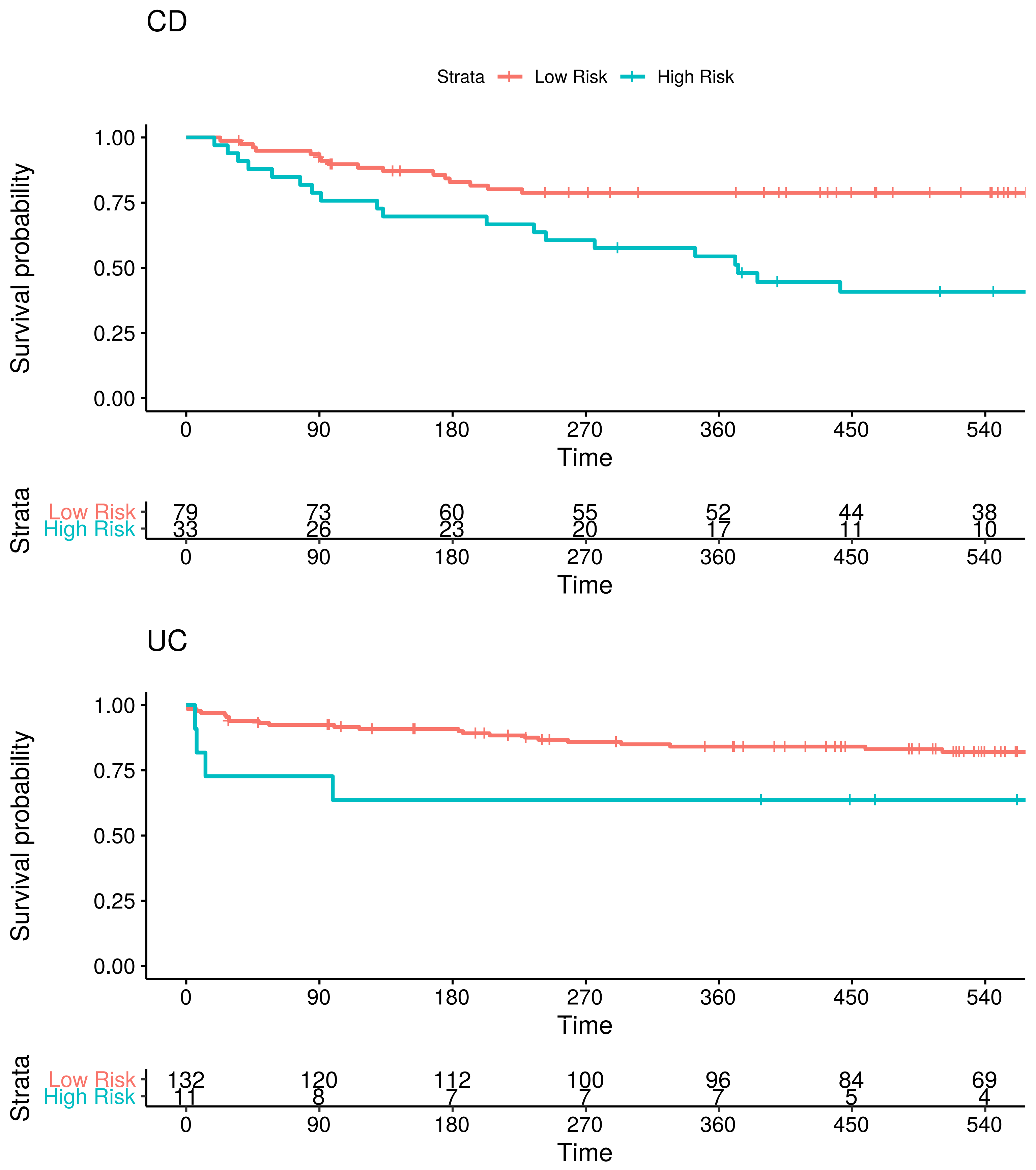

### Fig S4

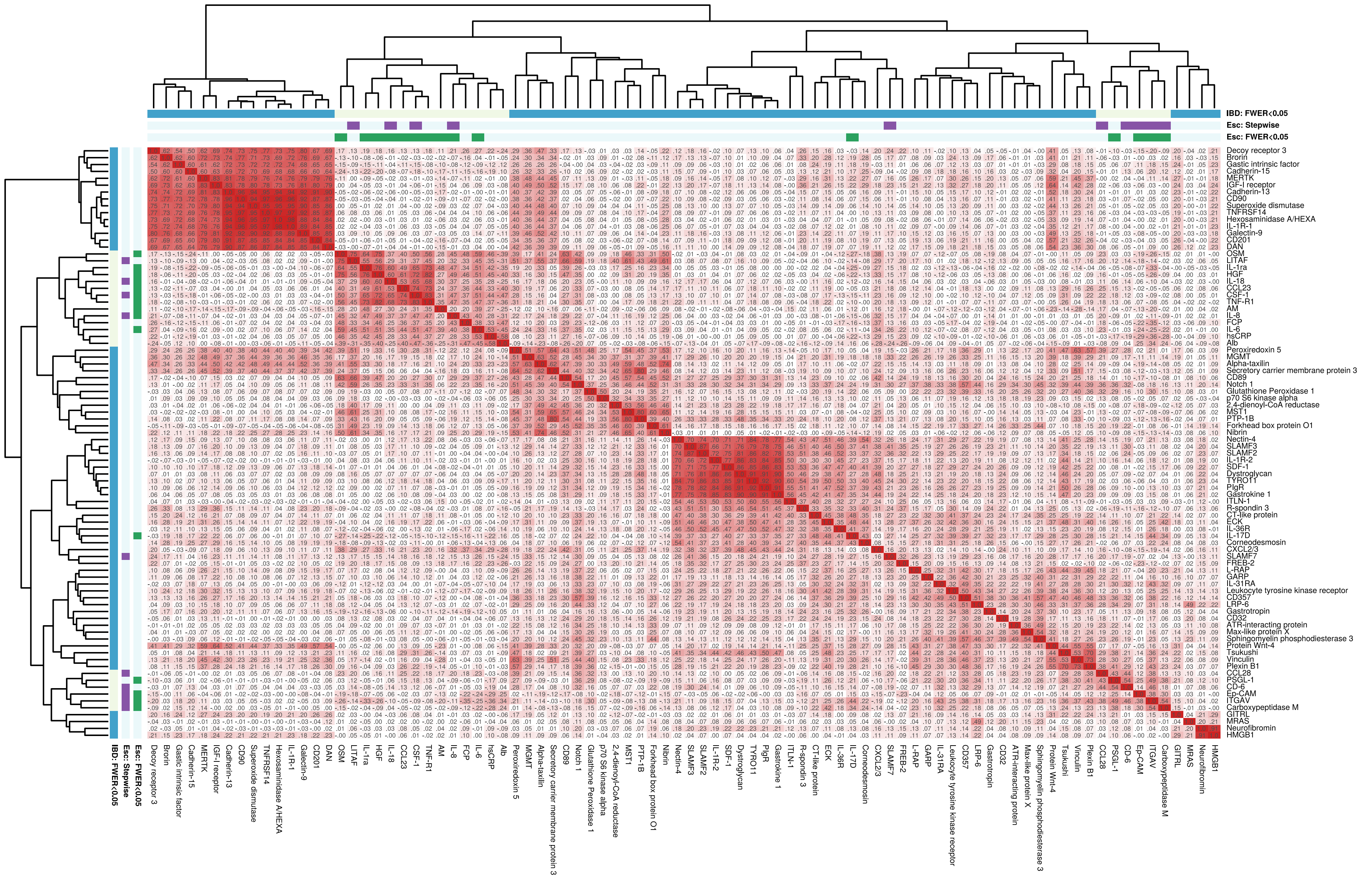

### Fig S5

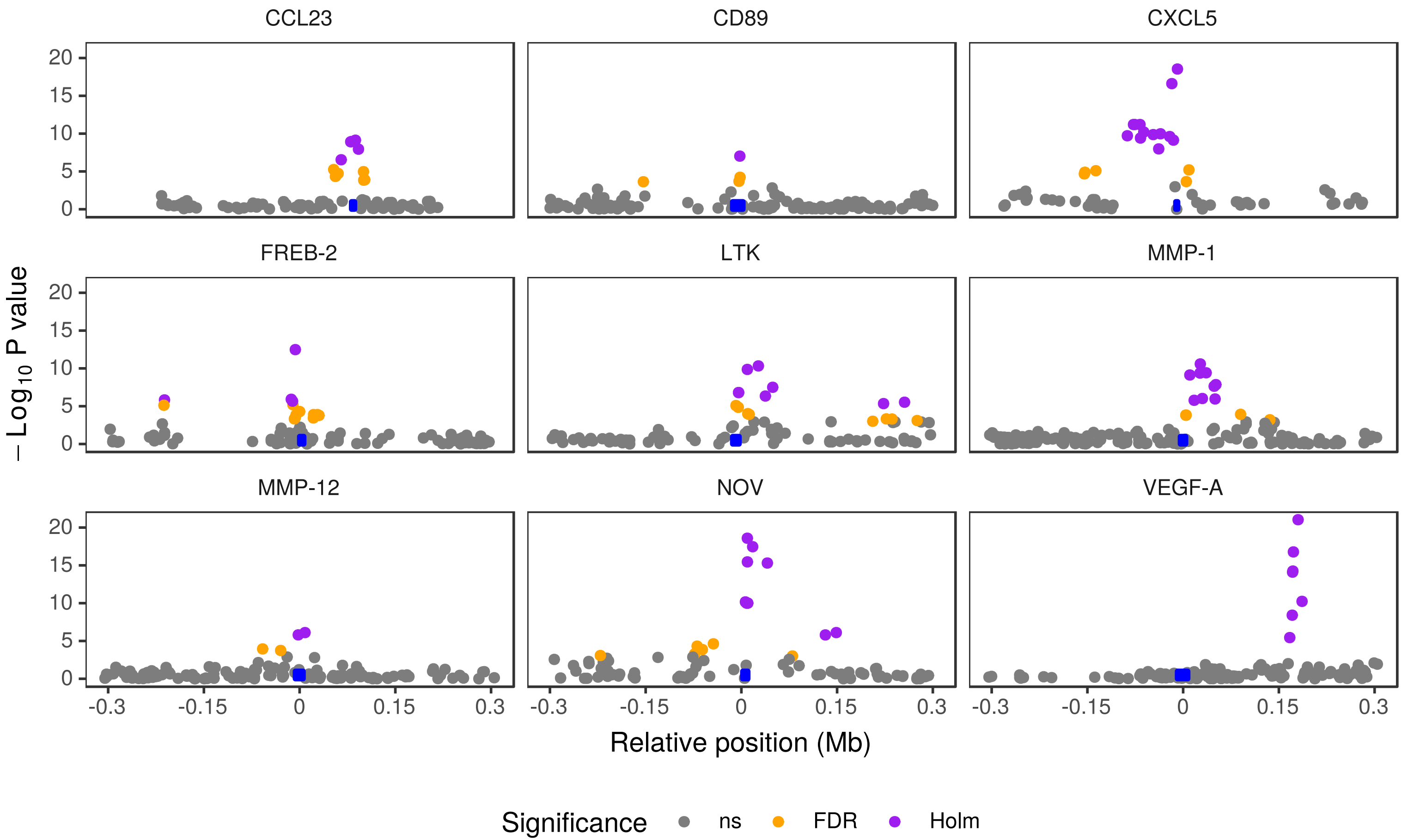

### Fig S6

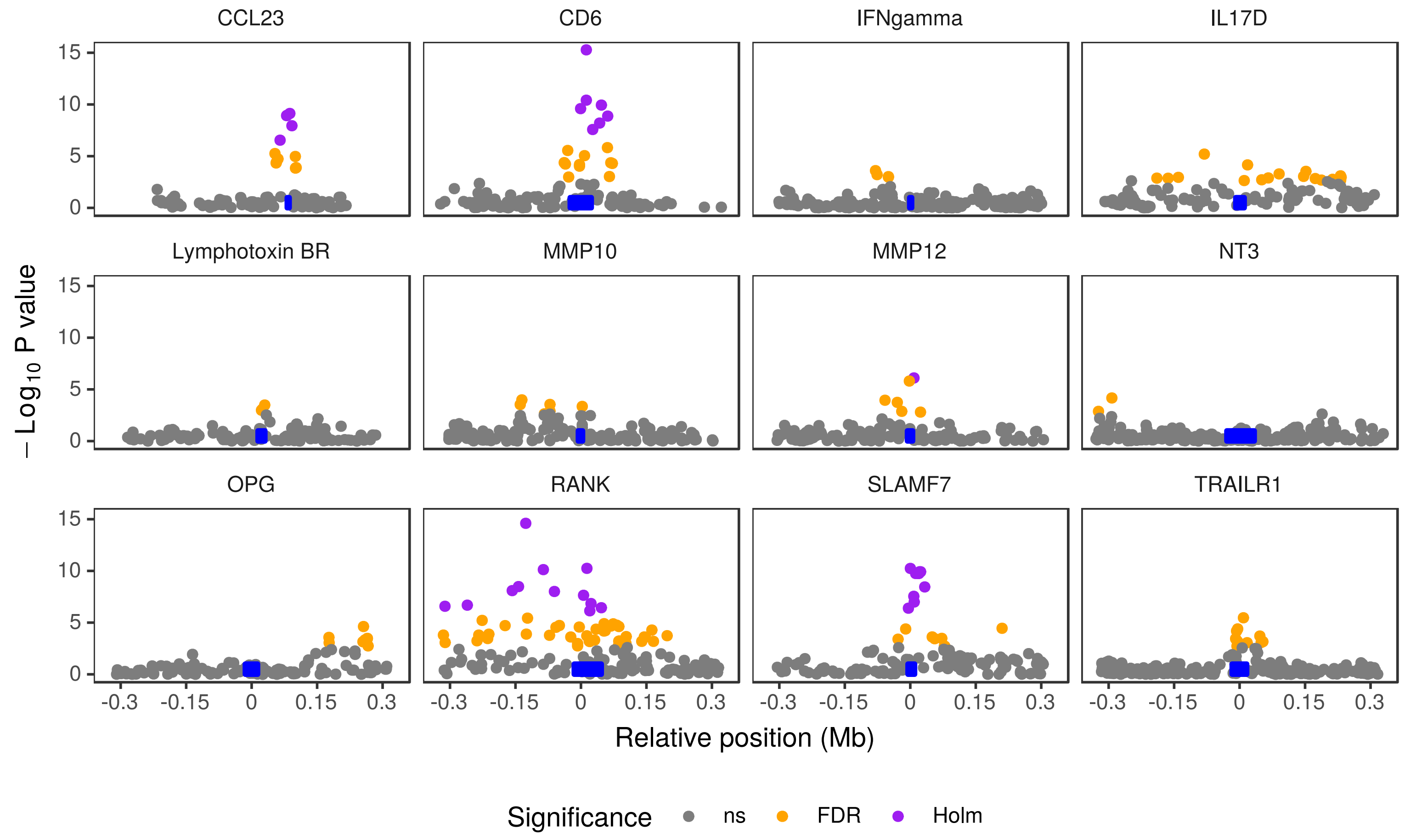

### Fig S7

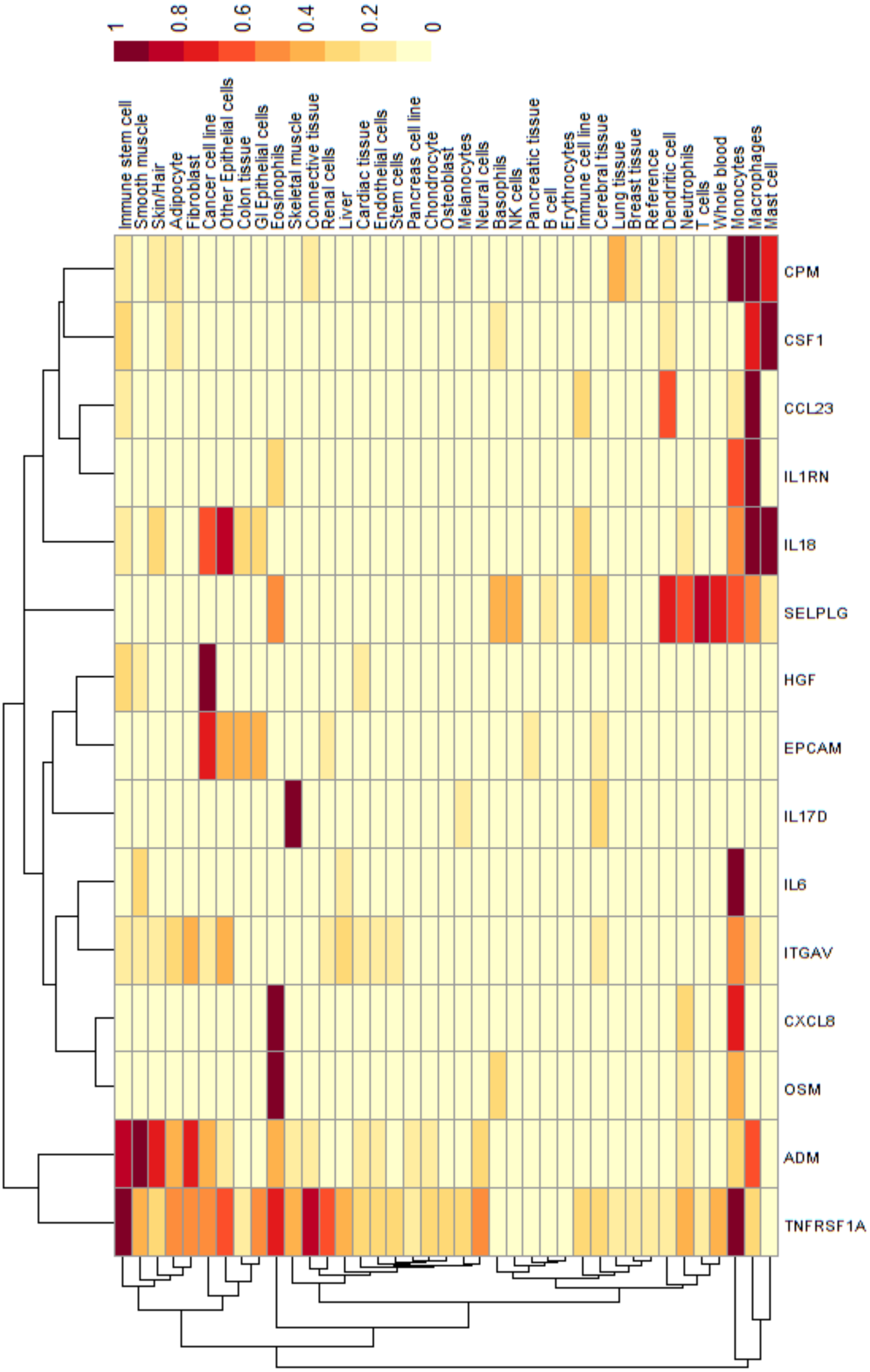
