## Supplementary tables for "Serum proteomic profiling at diagnosis predicts clinical course, and need for intensification of treatment in inflammatory bowel disease"

**Supplementary Table 1:** Percentage exposure of IBD patients to IBD-related medications for any indication during 30 days and 1 year prior to recruitment. Both exposures to Infliximab occurred >10 months prior to recruitment (ankylosing spondylitis)

|  |  | UC (143) | | CD (112) | | IBDU (23) | | Total | |
| --- | --- | --- | --- | --- | --- | --- | --- | --- | --- |
| Drug | Route/  Dose | <30 days | <1year | <30 days | <1year | <30 days | <1year | <30 days | <1year |
| Sulfasalazine | | 0 | 0 | 0.9 | 0.9 | 0 | 0 | 0.4 | 0.4 |
| 5-ASA | PO | 10.5 | 11.2 | 1.8 | 3.6 | 4.3 | 4.3 | 6.5 | 7.6 |
|  | PR | 7.7 | 12.6 | 0.9 | 0.9 | 4.3 | 4.3 | 4.7 | 7.2 |
|  | Total | 15.4 | 20.3 | 2.7 | 4.5 | 8.7 | 8.7 | 9.7 | 12.9 |
| Steroid | PR | 1.4 | 5.6 | 0 | 0 | 0 | 4.3 | 0.7 | 3.2 |
|  | IV | 4.9 | 5.6 | 6.3 | 8 | 8.7 | 8.7 | 5.8 | 6.8 |
|  | PO  High | 4.2 | 5.6 | 3.6 | 5.4 | 4.3 | 8.7 | 4 | 5.8 |
|  | PO  Med | 2.8 | 3.5 | 0.9 | 3.6 | 0 | 0 | 1.8 | 3.2 |
|  | PO  Low | 0 | 0.7 | 0 | 1.8 | 0 | 0 | 0 | 1.1 |
| AZA/6MP | | 0.7 | 1.4 | 1.8 | 1.8 | 0 | 0 | 1.1 | 1.4 |
| Ciclosporine | | 0 | 0.7 | 0.9 | 0.9 | 0 | 0 | 0.4 | 0.7 |
| Anti-TNF | | 0 | 0.7 | 0 | 0.9 | 0 | 0 | 0 | 0.7 |
| **Any Drug** | | **22.4** | **28.7** | **15.2** | **18.8** | **17.4** | **21.7** | **19.1** | **24.1** |

**Supplementary Table 2:** List of proteins profiled using the proximity extension assay in this study. * denotes exclusion as <50% of samples measured over the limit of detection (LOD).

| **UNIPROT** | **Samples > LOD (%)** | **Protein name** | **Gene Symbol** |
| --- | --- | --- | --- |
| Q16698 | 97.7 | 2,4-dienoyl-CoA reductase | DECR1 |
| P17050 | 72.4 | a-N-acetylgalactosaminidase | NAGA |
| P00813 | 97.2 | ADA | ADA |
| O00253 | 96.7 | AGRP | AGRP |
| P35475 | 98.1 | alpha-L-Iduronidase | IDUA |
| P40222 | 96.9 | Alpha-taxilin | TXLNA |
| P35318 | 96.7 | AM, Adrenomedullin | ADM |
| Q9BXJ7 | 96.5 | Amnionless | AMN |
| P04083 | 98.1 | ANXA1 | ANXA1 |
| P15514 | 96.5 | AR | AREG |
| Q8WXE1 | 63.0 | ATR-interacting protein | ATRIP |
| O15169 | 92.4 | AXIN1 | AXIN1 |
| Q9Y275 | 99.5 | BAFF | TNFSF13B |
| P23560 | 84.9 | BDNF | BDNF |
| P01138 | 87.0 | beta-NGF | NGF |
| P22004 | 98.1 | BMP-6 | BMP6 |
| Q9UK05 | 98.1 | BMP-9 | GDF2 |
| Q9BWV1 | 98.1 | BOC | BOC |
| Q2TAL6 | 96.9 | Brorin | VWC2 |
| Q7Z6A9 | 67.7 | BTLA | BTLA |
| Q8WXI7 | 96.7 | CA125 | MUC16 |
| P55290 | 96.9 | Cadherin-13 | CDH13 |
| P55291 | 96.9 | Cadherin-15 | CDH15 |
| Q16790 | 96.2 | CAIX | CA9 |
| P31949 | 96.9 | Calgizzarin | S100A11 |
| P35218 | 94.1 | Carbonic anhydrase 5A | CA5A |
| P14384 | 98.1 | Carboxypeptidase M | CPM |
| Q14790 | 95.3 | CASP-8 | CASP8 |
| P09668 | 97.7 | Cathepsin H | CTSH |
| P51671 | 99.4 | CCL11 | CCL11 |
| Q99731 | 99.4 | CCL19 | CCL19 |
| P78556 | 96.7 | CCL20 | CCL20 |
| P55773 | 99.5 | CCL23 | CCL23 |
| O15444 | 99.3 | CCL25 | CCL25 |
| Q9NRJ3 | 84.9 | CCL28 | CCL28 |
| P10147 | 96.5 | CCL3 | CCL3 |
| P13236 | 99.5 | CCL4 | CCL4 |
| P30203 | 96.5 | CD-6 | CD6 |
| O95971 | 98.1 | CD160 | CD160 |
| Q9UNN8 | 96.9 | CD201 | PROCR |
| P06734 | 98.1 | CD23 | FCER2 |
| Q9BZW8 | 99.5 | CD244 | CD244 |
| P32970 | 97.2 | CD27 Ligand | CD70 |
| P31994 | 98.1 | CD32 | FCGR2B |
| Q9Y5U5 | 53.5 | CD357 | TNFRSF18 |
| P01730 | 98.1 | CD4 | CD4 |
| P25942 | 99.3 | CD40 | CD40 |
| P29965 | 99.1 | CD40L | CD40LG |
| P06127 | 96.7 | CD5 | CD5 |
| Q07108 | 99.5 | CD69 | CD69 |
| P09564 | 97.0 | CD7 | CD7 |
| P24071 | 99.0 | CD89 | FCAR |
| P04216 | 96.9 | CD90 | THY1 |
| Q9H5V8 | 96.7 | CDCP1 | CDCP1 |
| P22223 | 96.7 | CDH3 | CDH3 |
| P38936 | 93.1 | CDKN1A | CDKN1A |
| P06731 | 55.2 | CEA | CEACAM5 |
| P01215 | 98.1 | CG alpha | CGA |
| Q15517 | 98.8 | Corneodesmosin | CDSN |
| P09603 | 99.5 | CSF-1 | CSF1 |
| P28325 | 99.5 | CST5 | CST5 |
| Q96IQ7 | 98.8 | CT-like protein | VSIG2 |
| P07711 | 99.5 | CTSL1 | CTSL |
| P78423 | 99.0 | CX3CL1 | CX3CL1 |
| P09341 | 99.5 | CXCL1 | CXCL1 |
| P02778 | 99.5 | CXCL10 | CXCL10 |
| O14625 | 99.3 | CXCL11 | CXCL11 |
| O43927 | 99.5 | CXCL13 | CXCL13 |
| P19876 | 99.0 | CXCL2/3 | CXCL3 |
| P42830 | 99.5 | CXCL5 | CXCL5 |
| P80162 | 99.5 | CXCL6 | CXCL6 |
| Q07325 | 99.5 | CXCL9 | CXCL9 |
| P41271 | 96.9 | DAN | NBL1 |
| Q9NNX6 | 98.1 | DC-SIGN | CD209 |
| P07585 | 98.1 | Decorin | DCN |
| O95407 | 96.9 | Decoy receptor 3 | TNFRSF6B |
| Q9BXN2 | 97.2 | Dectin-1 | CLEC7A |
| O94907 | 99.5 | Dkk-1 | DKK1 |
| Q8NFT8 | 99.3 | DNER | DNER |
| Q14118 | 98.9 | Dystroglycan | DAG1 |
| Q13541 | 96.0 | E4-BP1 | EIF4EBP1 |
| P29317 | 99.0 | ECK | EPHA2 |
| P12724 | 99.5 | ECP | RNASE3 |
| O43854 | 75.7 | EDIL3 | EDIL3 |
| P01133 | 99.3 | EGF | EGF |
| P00533 | 99.5 | EGFR | EGFR |
| P23588 | 93.2 | eIF-4B | EIF4B |
| P35613 | 99.5 | EMMPRIN, Basigin (Ok Blood Group) | BSG |
| P80511 | 96.7 | EN-RAGE | S100A12 |
| P16422 | 99.5 | Ep-CAM | EPCAM |
| O15197 | 98.1 | EphB6 | EPHB6 |
| P04626 | 99.5 | ErbB2/HER2 | ERBB2 |
| P21860 | 99.5 | ErbB3/HER3 | ERBB3 |
| Q15303 | 99.5 | ErbB4/HER4 | ERBB4 |
| Q9NQ30 | 99.5 | ESM-1 | ESM1 |
| P15311 | 99.3 | EZR | EZR |
| Q13158 | 96.7 | FADD | FADD |
| P25445 | 99.5 | FAS | FAS |
| P48023 | 96.5 | FasL | FASLG |
| O95750 | 99.5 | FGF-19 | FGF19 |
| Q9NSA1 | 96.5 | FGF-21 | FGF21 |
| Q9GZV9 | 54.9 | FGF-23 | FGF23 |
| P12034 | 91.5 | FGF-5 | FGF5 |
| O95633 | 98.1 | FLRG | FSTL3 |
| P49771 | 99.5 | Flt3L | FLT3LG |
| Q12778 | 91.3 | Forkhead box protein O1 | FOXO1 |
| P15328 | 96.7 | FR-alpha, Folate receptor alpha | FOLR1 |
| Q6BAA4 | 98.8 | FREB-2 | FCRLB |
| P19883 | 99.5 | FS | FST |
| Q12841 | 98.1 | FSTL1 | FSTL1 |
| P09958 | 99.5 | FUR | FURIN |
| P22466 | 99.5 | GAL | GAL |
| P09382 | 98.1 | Galectin-1 | LGALS1 |
| P05162 | 96.5 | Galectin-2 | LGALS2 |
| P56470 | 98.1 | Galectin-4 | LGALS4 |
| O00182 | 96.9 | Galectin-9 | LGALS9 |
| Q14392 | 60.1 | GARP | LRRC32 |
| P26992 | 98.1 | Gas6 | CNTFR |
| P27352 | 96.9 | Gastic intrinsic factor | GIF |
| Q9NS71 | 99.0 | Gastrokine 1 | GKN1 |
| P51161 | 81.8 | Gastrotropin | FABP6 |
| P39905 | 94.8 | GDNF | GDNF |
| O60609 | 98.1 | GFR alpha-3 | GFRA3 |
| P01241 | 98.8 | GH | GH1 |
| Q9UNG2 | 61.5 | GITRL | TNFSF18 |
| P35754 | 98.1 | Glutaredoxin 1 | GLRX |
| P07203 | 56.3 | Glutathione Peroxidase 1 | GPX1 |
| P35052 | 96.9 | Glypican-1 | GPC1 |
| P40189 | 91.8 | gp130 | IL6ST |
| Q14956 | 98.1 | GPNMB | GPNMB |
| P22749 | 98.1 | Granulysin | GNLY |
| P10144 | 97.7 | Granzyme B | GZMB |
| Q99075 | 99.5 | HB-EGF | HBEGF |
| Q14508 | 99.5 | HE4 | WFDC2 |
| P06865 | 96.9 | Hexosaminidase A/HEXA | HEXA |
| P14210 | 99.5 | HGF | HGF |
| P09429 | 80.7 | HMGB1 | HMGB1 |
| P09601 | 96.9 | HO-1 | HMOX1 |
| P04792 | 99.5 | HSP-27 | HSPB1 |
| Q9UJM8 | 98.1 | Hydroxyacid oxidase 1 | HAO1 |
| Q14773 | 96.9 | ICAM-4 | ICAM4 |
| O75144 | 96.7 | ICOSLG | ICOSLG |
| P01579 | 81.9 | IFN-gamma | IFNG |
| P15260 | 98.1 | IFN-gamma R1 | IFNGR1 |
| P38484 | 96.9 | IFN-gamma-R2 | IFNGR2 |
| P08069 | 96.9 | IGF-I receptor | IGF1R |
| P22301 | 95.5 | IL-10 | IL10 |
| Q08334 | 99.5 | IL-10RB | IL10RB |
| P29460 | 99.5 | IL-12 | IL12B |
| P29460 | 96.7 | IL-12B | IL12B |
| P78552 | 98.1 | IL-13 R alpha 1 | IL13RA1 |
| Q14005 | 96.7 | IL-16 | IL16 |
| Q96F46 | 98.1 | IL-17 RA | IL17RA |
| Q8NAC3 | 98.1 | IL-17 RC | IL17RC |
| Q16552 | 83.7 | IL-17A | IL17A |
| Q9P0M4 | 90.0 | IL-17C | IL17C |
| Q8TAD2 | 98.1 | IL-17D | IL17D |
| Q9NRM6 | 96.7 | IL-17RB | IL17RB |
| Q14116 | 99.5 | IL-18 | IL18 |
| Q13478 | 99.5 | IL-18R1 | IL18R1 |
| P14778 | 96.9 | IL-1R-1 | IL1R1 |
| P27930 | 99.0 | IL-1R-2 | IL1R2 |
| P18510 | 99.3 | IL-1ra | IL1RN |
| Q8NEV9 | 96.7 | IL-27 | IL27 |
| P26951 | 98.1 | IL-3 R alpha | IL3RA |
| Q8NI17 | 78.0 | IL-31RA | IL31RA |
| Q9HB29 | 99.0 | IL-36R | IL1RL2 |
| P24394 | 98.1 | IL-4 R alpha | IL4R |
| P05231 | 96.7 | IL-6 | IL6 |
| P13232 | 96.7 | IL-7 | IL7 |
| P10145 | 99.5 | IL-8 | CXCL8 |
| Q8NHJ6 | 96.4 | ILT-3 | LILRB4 |
| P05107 | 98.1 | Integrin beta 2 | ITGB2 |
| P56199 | 99.5 | ITGA1 | ITGA1 |
| P06756 | 98.1 | ITGAV | ITGAV |
| Q8WWA0 | 99.0 | ITLN-1 | ITLN1 |
| Q9UBX7 | 99.5 | K11 | KLK11 |
| Q92876 | 99.5 | KLK6 | KLK6 |
| Q6P179 | 73.8 | L-RAP | ERAP2 |
| P11279 | 98.1 | LAMP1 | LAMP1 |
| P13473 | 98.1 | LAMP2 | LAMP2 |
| Q9UJ71 | 97.4 | Langerin | CD207 |
| P01137 | 99.5 | LAP TGF-beta | TGFB1 |
| P01130 | 98.1 | LDL R | LDLR |
| P41159 | 94.3 | LEP | LEP |
| P29376 | 60.4 | Leukocyte tyrosine kinase receptor | LTK |
| P42702 | 75.0 | LIF-R | LIFR |
| Q99732 | 96.7 | LITAF | LITAF |
| P78380 | 99.5 | LOX-1 | OLR1 |
| O75581 | 78.8 | LRP-6 | LRP6 |
| P47992 | 98.1 | Lymphotactin | XCL1 |
| P36941 | 98.1 | Lymphotoxin beta R,  TNFReceptor Superfamily Member 3 | LTBR |
| P07948 | 95.7 | LYN | LYN |
| O43895 | 99.3 | mAmP | XPNPEP2 |
| Q9UEW3 | 98.1 | MARCO | MARCO |
| Q9UH92 | 58.3 | Max-like protein X | MLX |
| P13500 | 99.5 | MCP-1 | CCL2 |
| P80075 | 99.5 | MCP-2 | CCL8 |
| P80098 | 94.8 | MCP-3 | CCL7 |
| Q99616 | 96.7 | MCP-4 | CCL13 |
| Q12866 | 96.9 | MERTK | MERTK |
| P16455 | 94.6 | MGMT | MGMT |
| Q16674 | 99.5 | MIA | MIA |
| Q29983 | 87.8 | MIC-A | MICA |
| P10147 | 96.5 | MIP-1 alpha | CCL3 |
| P21741 | 94.8 | MK | MDK |
| P03956 | 99.1 | MMP-1 | MMP1 |
| P09238 | 99.5 | MMP-10 | MMP10 |
| P39900 | 99.5 | MMP-12 | MMP12 |
| P09237 | 99.5 | MMP-7 | MMP7 |
| P08253 | 98.1 | MMP2 | MMP2 |
| O14807 | 68.1 | MRAS | MRAS |
| Q13043 | 96.4 | MST1 | STK4 |
| P19022 | 98.1 | N-Cadherin | CDH2 |
| Q96NY8 | 99.0 | Nectin-4 | NECTIN4 |
| Q9Y6K9 | 96.7 | NEMO | IKBKG |
| P21359 | 99.0 | Neurofibromin | NF1 |
| O60934 | 94.1 | Nibrin | NBN |
| O76036 | 85.1 | NK-p46 | NCR1 |
| P46531 | 97.0 | Notch 1 | NOTCH1 |
| P48745 | 98.1 | NOV | NOV |
| P20783 | 88.7 | NT-3 | NTF3 |
| P16860 | 79.2 | NT-pro-BNP | NPPB |
| O75354 | 96.9 | NTPDase 6 | ENTPD6 |
| Q16288 | 99.5 | NTRK3 | NTRK3 |
| O00300 | 99.5 | OPG | TNFRSF11B |
| P13725 | 96.7 | OSM | OSM |
| P46527 | 91.5 | p27/Kip1/CDKN1B | CDKN1B |
| P23443 | 77.8 | p70 S6 kinase alpha | RPS6KB1 |
| Q13219 | 99.5 | PAPP-A | PAPPA |
| P25116 | 96.7 | PAR-1 | F2R |
| Q99497 | 99.5 | PARK7 | PARK7 |
| Q15116 | 70.1 | PD-1 | PDCD1 |
| Q9NZQ7 | 82.3 | PD-L1 | CD274 |
| Q9BQ51 | 98.1 | PD-L2 | PDCD1LG2 |
| P01127 | 99.5 | PDGF subunit B | PDGFB |
| P30044 | 95.3 | Peroxiredoxin 5 | PRDX5 |
| P49763 | 99.5 | PIGF, Placental growth factor | PGF |
| P01833 | 99.0 | PIgR | PIGR |
| O43157 | 91.7 | Plexin B1 | PLXNB1 |
| P01236 | 99.5 | PRL | PRL |
| P56705 | 89.4 | Protein Wnt-4 | WNT4 |
| Q16651 | 99.5 | PRSS8 | PRSS8 |
| Q14242 | 96.7 | PSGL-1 | SELPLG |
| P18031 | 99.0 | PTP-1B | PTPN1 |
| Q9Y2R2 | 96.7 | PTPN22 | PTPN22 |
| P26022 | 96.4 | PTX3 | PTX3 |
| Q9BXY4 | 99.0 | R-spondin 3 | RSPO3 |
| Q15109 | 99.5 | RAGE | AGER |
| Q9Y6Q6 | 98.1 | RANK | TNFRSF11A |
| Q9BYZ8 | 99.3 | REG-4 | REG4 |
| P00797 | 99.5 | REN | REN |
| P07949 | 98.1 | Ret | RET |
| Q14108 | 98.1 | SCARB2 | SCARB2 |
| P21583 | 99.5 | SCF | KITLG |
| P48061 | 98.9 | SDF-1 | CXCL12 |
| O14828 | 96.4 | Secretory carrier membrane protein 3 | SCAMP3 |
| Q9BQR3 | 98.1 | Serine protease 27 | PRSS27 |
| P78324 | 98.1 | SIRP alpha | SIRPA |
| Q8IXJ6 | 96.7 | SIRT2 | SIRT2 |
| Q13291 | 95.1 | SLAMF1 | SLAMF1 |
| P09326 | 99.0 | SLAMF2 | CD48 |
| Q9HBG7 | 99.0 | SLAMF3 | LY9 |
| Q9UIB8 | 98.1 | SLAMF5 | CD84 |
| Q9NQ25 | 90.6 | SLAMF7 | SLAMF7 |
| Q99717 | 96.9 | SMAD 5 | SMAD5 |
| Q9NY59 | 57.3 | Sphingomyelin phosphodiesterase 3 | SMPD3 |
| Q9HCB6 | 99.5 | SPON1 | SPON1 |
| P50225 | 88.5 | ST1A1 | SULT1A1 |
| O95630 | 96.7 | STAMPB | STAMBP |
| P04179 | 96.9 | Superoxide dismutase | SOD2 |
| P78536 | 79.5 | TACE | ADAM17 |
| O14836 | 98.1 | TACI | TNFRSF13B |
| P13726 | 99.5 | TF, tissue Factor | F3 |
| Q07654 | 98.1 | TFF3 | TFF3 |
| P48307 | 98.1 | TFPI-2 | TFPI2 |
| P01135 | 96.5 | TGFA | TGFA |
| P40225 | 96.5 | THPO | THPO |
| Q02763 | 99.5 | TIE-2 | TEK |
| Q96D42 | 98.4 | TIM | HAVCR1 |
| Q15399 | 85.8 | TLR1 | TLR1 |
| O60603 | 97.7 | TLR2 | TLR2 |
| O15455 | 98.1 | TLR3 | TLR3 |
| P07204 | 99.5 | TM | THBD |
| P19438 | 99.5 | TNF-R1 | TNFRSF1A |
| P01374 | 96.7 | TNFB | LTA |
| Q92956 | 96.9 | TNFRSF14 | TNFRSF14 |
| O75509 | 98.1 | TNFRSF21 | TNFRSF21 |
| P43489 | 96.7 | TNFRSF4 | TNFRSF4 |
| P26842 | 98.1 | TNFRSF7 | CD27 |
| Q07011 | 99.5 | TNFRSF9 | TNFRSF9 |
| O43557 | 96.9 | TNFSF14 | TNFSF14 |
| P50591 | 99.5 | TRAIL | TNFSF10 |
| O00220 | 98.1 | TRAIL R1 | TNFRSF10A |
| O14763 | 96.7 | TRAIL-R2 | TNFRSF10B |
| O14788 | 96.4 | TRANCE | TNFSF11 |
| P21980 | 98.1 | Transglutaminase 2 | TGM2 |
| Q9NP99 | 97.9 | TREM-1 | TREM1 |
| Q9NZC2 | 98.1 | TREM-2 | TREM2 |
| Q5T2D2 | 98.1 | TREML2 | TREML2 |
| Q9NS68 | 98.1 | TROY | TNFRSF19 |
| Q8WUA8 | 99.0 | Tsukushi | TSKU |
| O43508 | 99.5 | TWEAK | TNFSF12 |
| P54760 | 99.0 | TYRO11 | EPHB4 |
| Q03405 | 99.5 | U-PAR | PLAUR |
| P00749 | 99.5 | uPA | PLAU |
| Q9UHF1 | 89.8 | VE-statin | EGFL7 |
| P15692 | 99.5 | VEGF-A | VEGFA |
| O43915 | 99.5 | VEGF-D | VEGFD |
| P35968 | 99.5 | VEGFR-2 | KDR |
| P08670 | 96.7 | VIM | VIM |
| P18206 | 66.8 | Vinculin | VCL |
| Q9Y5W5 | 98.1 | WIF1 | WIF1 |
| P14061 | 24.7* | 17-beta-HSD 1 | HSD17B1 |
| Q15025 | 48.6* | ABIN-1 | TNIP1 |
| P55263 | 19.4* | Adenosine Kinase | ADK |
| P31749 | 23.8* | AKT Serine/Threonine Kinase 1 | AKT1 |
| O15144 | 23.8* | Actin Related Protein 2/3 Complex Subunit 2 | ARPC2 |
| Q5T4W7 | 16.1* | Artemin | ARTN |
| O43918 | 9.5* | Autoimmune Regulator | AIRE |
| Q9NY97 | 9.9* | B3GNT2 | B3GNT2 |
| Q14451 | 33.2* | B47 | GRB7 |
| Q92934 | 5.2* | Bcl2 Antagonist Of Cell Death | BAD |
| O95999 | 8.0* | BCL10 | BCL10 |
| P41182 | 26.2* | BCL6 | BCL6 |
| Q86UU0 | 11.3* | BCL9-2 | BCL9L |
| O75626 | 10.6* | BLIMP1 | PRDM1 |
| P36894 | 5.9* | BMPR-IA | BMPR1A |
| P16860 | 13.4* | BNP | NPPB |
| Q9NXR7 | 17.4* | BRCA1-A complex subunit BRE | BABAM2 |
| Q13882 | 24.1* | BRK | PTK6 |
| P05937 | 25.7* | Calbindin | CALB1 |
| P05109 | 5.6* | Calgranulin-A | S100A8 |
| P06702 | 19.8* | Calgranulin-B | S100A9 |
| Q13555 | 9.2* | CaMK-II subunit gamma | CAMK2G |
| Q9BXL7 | 4.1* | CARD11 | CARD11 |
| Q9H257 | 24.7* | CARD9s | CARD9 |
| P57735 | 11.3* | CATX-8 | RAB25 |
| Q03135 | 37.5* | Caveolin 1 | CAV1 |
| P22362 | 33.7* | CCL1 | CCL1 |
| P51684 | 10.9* | CCR-6 | CCR6 |
| Q16832 | 17.5* | CD167b | DDR2 |
| Q00610 | 12.0* | CLH-17 | CLTC |
| P26441 | 5.9* | CNTF | CNTF |
| P13073 | 17.0* | COX IV-1 | COX4I1 |
| O14490 | 38.9* | DAP-1 | DLGAP1 |
| Q5S007 | 9.0* | Dardarin | LRRK2 |
| O14640 | 31.3* | Dishevelled-1 | DVL1 |
| Q13422 | 18.2* | DNA-binding protein Ikaros | IKZF1 |
| P26358 | 34.4* | Dnmt1 | DNMT1 |
| Q9UBC3 | 11.5* | Dnmt3b | DNMT3B |
| Q9UBSB | 36.1* | E3 ubiquitin-protein ligase RNF14 | RNF14 |
| Q9NYQ7 | 21.7* | EGF-like-protein 1 | CELSR3 |
| P21709 | 24.8* | EPHA1 | EPHA1 |
| P01588 | 45.5* | Erythropoietin | EPO |
| P26885 | 13.4* | FKBP-13 | FKBP2 |
| Q9NZU1 | 22.6* | FLRT1 | FLRT1 |
| P15407 | 25.3* | FRA-1 | FOSL1 |
| Q495W5 | 31.1* | Fucosyltransferase 11 | FUT11 |
| P04141 | 15.3* | GM-CSF | CSF2 |
| P60983 | 7.8* | GMFB | GMFB |
| P49841 | 7.5* | GSK3B | GSK3B |
| P41235 | 13.4* | HNF-4-alpha | HNF4A |
| P49639 | 3.5* | Homeobox protein Hox-A1 | HOXA1 |
| Q86Z02 | 13.7* | Homeodomain-interacting protein kinase 1 | HIPK1 |
| Q9UMF0 | 6.3* | ICAM-5 | ICAM5 |
| P17181 | 27.1* | IFN-R-1 | IFNAR1 |
| P48551 | 12.8* | IFN-R-2 | IFNAR2 |
| P01583 | 5.0* | IL-1 alpha | IL1A |
| P01584 | 4.3* | IL-1-beta | IL1B |
| Q13651 | 27.8* | IL10-RA | IL10RA |
| Q99665 | 1.9* | IL-12RB2 | IL12RB2 |
| P35225 | 11.3* | IL-13 | IL13 |
| Q13261 | 34.4* | IL-15RA | IL15RA |
| Q9UHF5 | 32.3* | IL-17B | IL17B |
| O95256 | 8.2* | IL-18RAcP | IL18RAP |
| Q9UHD0 | 4.5* | IL-19 | IL19 |
| P60568 | 16.0* | IL-2 | IL2 |
| Q9NYY1 | 21.2* | IL-20 | IL20 |
| Q9UHF4 | 10.6* | IL-20RA | IL20RA |
| Q9GZX6 | 21.2* | IL-22 | IL22 |
| Q8N6P7 | 33.2* | IL-22 RA1 | IL22RA1 |
| Q5VWK5 | 13.7* | IL-23R | IL23R |
| Q13007 | 24.3* | IL-24 | IL24 |
| Q8IU54 | 20.8* | IL-29 | IFNL1 |
| P14784 | 28.6* | IL-2RB | IL2RB |
| O95760 | 7.5* | IL-33 | IL33 |
| Q6ZMJ4 | 20.7* | IL-34 | IL34 |
| Q9NZH7 | 20.8* | IL-36 beta | IL36B |
| P05112 | 8.2* | IL-4 | IL4 |
| P05113 | 26.6* | IL-5 | IL5 |
| P14735 | 4.2* | Insulysin | IDE |
| P01562 | 19.8* | Interferon alpha-D | IFNA1 |
| P46940 | 32.8* | IQGAP1 | IQGAP1 |
| Q9UIQ6 | 19.6* | IRAP | LNPEP |
| Q13568 | 17.0* | IRF-5 | IRF5 |
| P10914 | 41.0* | IRF1 | IRF1 |
| Q02556 | 25.2* | IRF8 | IRF8 |
| P35568 | 25.2* | IRS1 | IRS1 |
| Q96A47 | 6.6* | ISL2 | ISL2 |
| Q9UKP3 | 4.5* | ITGB1BP2 | ITGB1BP2 |
| P68036 | 9.7* | L-UBC | UBE2L3 |
| Q8IV20 | 2.3* | LACC1 | LACC1 |
| P09960 | 48.4* | Leukotriene A4 Hydrolase | LTA4H |
| O60711 | 28.3* | Leupaxin | LPXN |
| P15018 | 13.9* | LIF | LIF |
| Q15831 | 25.7* | Serine/Threonine Kinase 11 | LKB1 |
| Q9HBW0 | 28.0* | Lysophosphatidic Acid Receptor 2 | LPAR2 |
| A1A4Y4 | 1.0* | LRG-47 | IRGM |
| Q9Y561 | 8.9* | LRP-12 | LRP12 |
| O00462 | 8.3* | Mannase | MANBA |
| P28482 | 9.0* | MAPK 1 | MAPK1 |
| O95819 | 9.7* | MEKKK 4 | MAP4K4 |
| Q29980 | 47.4* | MIC-B | MICB |
| Q04912 | 11.1* | MSP receptor | MST1R |
| Q99836 | 17.2* | MYD88 | MYD88 |
| P46934 | 20.8* | NEDD-4 | NEDD4 |
| Q495T6 | 24.0* | Neprilysin-2 | MMEL1 |
| Q15784 | 24.1* | NeuroD2 | NEUROD2 |
| O14931 | 26.7* | NKp30 | NCR3 |
| Q04721 | 9.0* | Notch-2 | NOTCH2 |
| Q99748 | 8.2* | Neurturin | NRTN |
| Q9NX02 | 13.7* | Neucleotide-binding site protein 1 | NLRP2 |
| Q01860 | 4.0* | Oct-3 | POU5F1 |
| Q9Y4L1 | 27.1* | ORP150 | HYOU1 |
| O60356 | 26.8* | P8 / NUPR1 | NUPR1 |
| Q05655 | 32.6* | PKC delta | PRKCD |
| P05771 | 22.9* | PKC-beta 1 | PRKCB |
| P05771 | 1.6* | PKC-beta 2 | PRKCB |
| Q01970 | 34.5* | PLC-beta-3 | PLCB3 |
| P53350 | 23.8* | PLK-1 | PLK1 |
| P11086 | 10.9* | PNMTase | PNMT |
| O15355 | 32.1* | PP2C-gamma | PPM1G |
| P01100 | 26.9* | Proto-oncogene c-Fos | FOS |
| Q92878 | 2.1* | RAD50 | RAD50 |
| Q8TDF6 | 41.3* | RASGRP4 | RASGRP4 |
| O43781 | 12.2* | REDK | DYRK3 |
| Q9P0U3 | 14.6* | SENP1 | SENP1 |
| Q6IA17 | 37.2* | SIGIRR | SIGIRR |
| Q15796  P84022 | 4.0* | SMAD 2-3 | SMAD2  SMAD3 |
| O15198 | 14.8* | SMAD9 | SMAD9 |
| Q7Z699 | 24.7* | Spred-1 | SPRED1 |
| Q7Z698 | 17.2* | Spred-2 | SPRED2 |
| Q9C004 | 2.8* | Spry-4 | SPRY4 |
| P12931 | 4.0* | SRC | SRC |
| O00204 | 37.8* | ST2B1 | SULT2B1 |
| P42224 | 1.0* | STAT-1 | STAT1 |
| P40763 | 5.7* | STAT-3 | STAT3 |
| Q14765 | 22.4* | STAT-4 | STAT4 |
| P42229 | 6.8* | STAT-5A | STAT5A |
| Q15750 | 17.5* | TAK1-binding protein 1 | TAB1 |
| P17706 | 37.2* | TCPTP | PTPN2 |
| P01375 | 13.2* | TNF | TNF |
| Q04206 | 24.5* | Transcription factor p65 | RELA |
| Q969D9 | 14.2* | TSLP | TSLP |
| Q9Y2W6 | 36.1* | Tudor and KH domain-containing protein | TDRKH |
| Q9BZM6 | 15.5* | ULBP-1 | ULBP1 |
| P49765 | 15.1* | VEGF-B | VEGFB |
| Q9ULJ6 | 11.1* | Zimp10 | ZMIZ1 |
| P17028 | 38.2* | ZNF24 | ZNF24 |

**Supplementary Table 3:** - Table of differentially expressed protein markers between inflammatory bowel disease (IBD) cases and controls in serum with age and sex as covariates. Log_2_ Fold-Change denotes mean IBD expression – mean control expression, with expression as normalised Ct values (absolute values < 0.01 are reported as 0). Proteins which lost significance when inflammatory covariates were included are marked with *.

| Protein | Log_2_FC | P Value | Holm P |
| --- | --- | --- | --- |
| MMP-12 | 0.87 | 1.32×10^-25^ | 4.14×10^-23^ |
| CXCL1 | 0.78 | 1.28×10^-19^ | 3.99×10^-17^ |
| IL-8 | 0.83 | 3.34×10^-19^ | 1.04×10^-16^ |
| OSM | 0.81 | 1.19×10^-18^ | 3.67×10^-16^ |
| IL-17A | 0.69 | 1.20×10^-18^ | 3.69×10^-16^ |
| CXCL9 | 0.84 | 5.68×10^-17^ | 1.75×10^-14^ |
| Granzyme B | 1.05 | 2.16×10^-15^ | 6.62×10^-13^ |
| MMP-10 | 0.65 | 1.20×10^-13^ | 3.68×10^-11^ |
| HGF | 0.49 | 4.58×10^-13^ | 1.40×10^-10^ |
| CXCL11 | 0.59 | 7.53×10^-13^ | 2.29×10^-10^ |
| CXCL2/3 | 0.79 | 5.63×10^-12^ | 1.71×10^-9^ |
| CCL20 | 0.63 | 6.87×10^-12^ | 2.07×10^-9^ |
| Gas6 | -0.18 | 1.70×10^-11^ | 5.12×10^-9^ |
| AR | 0.47 | 4.59×10^-11^ | 1.38×10^-8^ |
| ITGAV | -0.27 | 6.01×10^-11^ | 1.80×10^-8^ |
| IL-6 | 0.80 | 8.17×10^-11^ | 2.43×10^-8^ |
| IL-17C | 0.38 | 8.03×10^-10^ | 2.39×10^-7^ |
| Decoy receptor 3 | 0.60 | 2.04×10^-9^ | 6.05×10^-7^ |
| IL-1RA | 0.47 | 2.05×10^-9^ | 6.05×10^-7^ |
| FR-alpha | -0.23 | 2.30×10^-9^ | 6.77×10^-7^ |
| TRANCE | -0.44 | 4.99×10^-9^ | 1.46×10^-6^ |
| NOV | -0.17 | 1.25×10^-8^ | 3.66×10^-6^ |
| CCL11 | 0.29 | 2.43×10^-8^ | 7.07×10^-6^ |
| SCF | -0.39 | 3.58×10^-8^ | 1.04×10^-5^ |
| TGFA | 0.42 | 3.98×10^-8^ | 1.15×10^-5^ |
| Galectin-1 | -0.16 | 4.39×10^-8^ | 1.27×10^-5^ |
| Carboxypeptidase M | -0.19 | 6.10×10^-8^ | 1.75×10^-5^ |
| IL-10 | 0.49 | 6.40×10^-8^ | 1.83×10^-5^ |
| IL-4 R alpha | 0.28 | 1.12×10^-7^ | 3.19×10^-5^ |
| FREB-2 | 0.27 | 2.24×10^-7^ | 6.37×10^-5^ |
| ICOSLG | -0.38 | 3.08×10^-7^ | 8.72×10^-5^ |
| IL-17D | -0.16 | 3.64×10^-7^ | 1.03×10^-4^ |
| PTPN22 | 0.43 | 3.71×10^-7^ | 1.04×10^-4^ |
| CSF-1 | 0.31 | 4.94×10^-7^ | 1.38×10^-4^ |
| IFN-gamma | 0.45 | 5.59×10^-7^ | 1.56×10^-4^ |
| GFR alpha-3 | -0.19 | 5.81×10^-7^ | 1.62×10^-4^ |
| TROY | -0.19 | 6.44×10^-7^ | 1.78×10^-4^ |
| REG-4 | 0.37 | 7.83×10^-7^ | 2.16×10^-4^ |
| IL-7 | 0.26 | 9.43×10^-7^ | 2.59×10^-4^ |
| LITAF | 0.41 | 1.18×10^-6^ | 3.23×10^-4^ |
| VEGF-A | 0.32 | 1.21×10^-6^ | 3.31×10^-4^ |
| DNER | -0.24 | 2.05×10^-6^ | 5.56×10^-4^ |
| Langerin* | -0.22 | 3.41×10^-6^ | 9.24×10^-4^ |
| gp130 | -0.21 | 4.92×10^-6^ | 1.33×10^-3^ |
| ITGA1 | -0.24 | 9.53×10^-6^ | 2.56×10^-3^ |
| TNFB* | -0.22 | 9.55×10^-6^ | 2.56×10^-3^ |
| CDH3 | -0.20 | 9.79×10^-6^ | 2.62×10^-3^ |
| MMP-1* | 0.41 | 1.29×10^-5^ | 3.44×10^-3^ |
| BOC | -0.16 | 1.48×10^-5^ | 3.93×10^-3^ |
| MCP-3 | 0.24 | 2.38×10^-5^ | 6.28×10^-3^ |
| CCL23 | 0.28 | 2.79×10^-5^ | 7.33×10^-3^ |
| ErbB2/HER2 | -0.20 | 3.39×10^-5^ | 8.89×10^-3^ |
| IL-36R* | -0.22 | 3.40×10^-5^ | 8.89×10^-3^ |
| Ret* | -0.20 | 3.81×10^-5^ | 9.90×10^-3^ |
| Nibrin | 0.30 | 5.19×10^-5^ | 1.34×10^-2^ |
| CXCL5* | 0.31 | 6.05×10^-5^ | 1.56×10^-2^ |
| CD89* | 0.22 | 6.47×10^-5^ | 1.66×10^-2^ |
| AM | 0.29 | 7.14×10^-5^ | 1.83×10^-2^ |
| Gastrotropin* | -0.18 | 7.62×10^-5^ | 1.94×10^-2^ |
| EGFR | -0.15 | 8.00×10^-5^ | 2.03×10^-2^ |
| BMP-9* | -0.23 | 1.10×10^-4^ | 2.77×10^-2^ |
| Leukocyte tyrosine kinase receptor* | -0.10 | 1.24×10^-4^ | 3.12×10^-2^ |
| Flt3L | -0.22 | 1.64×10^-4^ | 4.12×10^-2^ |
| TNF-R1* | 0.20 | 1.70×10^-4^ | 4.24×10^-2^ |
| EMMPRIN | -0.18 | 1.71×10^-4^ | 4.26×10^-2^ |
| IL-27 | 0.17 | 1.78×10^-4^ | 4.43×10^-2^ |
| ILT-3 | 0.16 | 2.34×10^-4^ | 0.06 |
| Ep-CAM | -0.33 | 2.64×10^-4^ | 0.07 |
| FUR | 0.17 | 3.26×10^-4^ | 0.08 |
| LAMP2 | -0.08 | 3.30×10^-4^ | 0.08 |
| mAmP | -0.39 | 3.73×10^-4^ | 0.09 |
| Alpha-taxilin | 0.23 | 3.82×10^-4^ | 0.09 |
| PTX3 | 0.26 | 3.88×10^-4^ | 0.09 |
| Corneodesmosin | -0.19 | 3.95×10^-4^ | 0.09 |
| TRAIL R1 | 0.16 | 4.02×10^-4^ | 0.10 |
| PSGL-1 | -0.08 | 4.16×10^-4^ | 0.10 |
| IL-18 | 0.20 | 4.28×10^-4^ | 0.10 |
| NT-3 | -0.14 | 4.63×10^-4^ | 0.11 |
| IL-17 RC | -0.19 | 4.67×10^-4^ | 0.11 |
| MCP-4 | -0.18 | 5.28×10^-4^ | 0.12 |
| LEP | -0.41 | 5.62×10^-4^ | 0.13 |
| GPNMB | -0.11 | 5.80×10^-4^ | 0.14 |
| MIA | -0.20 | 8.65×10^-4^ | 0.20 |
| Tsukushi | -0.32 | 8.86×10^-4^ | 0.20 |
| LDL R | -0.09 | 8.93×10^-4^ | 0.20 |
| Plexin B1 | -0.36 | 9.11×10^-4^ | 0.21 |
| SLAMF7 | 0.17 | 9.25×10^-4^ | 0.21 |
| RAGE | -0.17 | 1.03×10^-3^ | 0.23 |
| Decorin | -0.09 | 1.11×10^-3^ | 0.25 |
| CCL19 | 0.27 | 1.19×10^-3^ | 0.27 |
| NTRK3 | -0.15 | 1.33×10^-3^ | 0.30 |
| TLR2 | 0.13 | 1.49×10^-3^ | 0.33 |
| SMAD 5 | -0.19 | 1.58×10^-3^ | 0.35 |
| TNFRSF21 | -0.09 | 1.62×10^-3^ | 0.36 |
| ECK | -0.13 | 1.63×10^-3^ | 0.36 |
| ECP | 0.36 | 1.81×10^-3^ | 0.39 |
| EDIL3 | -0.16 | 2.04×10^-3^ | 0.44 |
| K11 | -0.13 | 2.28×10^-3^ | 0.49 |
| OPG | 0.16 | 2.31×10^-3^ | 0.50 |
| TNFRSF7 | 0.12 | 2.46×10^-3^ | 0.53 |
| TREM-1 | 0.13 | 2.94×10^-3^ | 0.63 |
| PDGF subunit B | 0.17 | 4.32×10^-3^ | 0.92 |
| Cadherin-15 | -0.27 | 4.47×10^-3^ | 0.94 |
| CXCL13 | 0.20 | 4.99×10^-3^ | 1.00 |
| IL-31RA | -0.09 | 5.12×10^-3^ | 1.00 |
| KLK6 | -0.14 | 5.17×10^-3^ | 1.00 |
| BAFF | 0.13 | 5.38×10^-3^ | 1.00 |
| Vinculin | -0.16 | 5.65×10^-3^ | 1.00 |
| TF | -0.11 | 6.39×10^-3^ | 1.00 |
| TREM-2 | -0.12 | 6.79×10^-3^ | 1.00 |
| NEMO | 0.18 | 7.36×10^-3^ | 1.00 |
| CXCL6 | 0.2 | 7.64×10^-3^ | 1.00 |
| CCL4 | 0.16 | 8.23×10^-3^ | 1.00 |
| PTP-1B | 0.21 | 8.32×10^-3^ | 1.00 |
| BMP-6 | 0.12 | 9.36×10^-3^ | 1.00 |
| Glutathione Peroxidase 1 | -0.10 | 9.38×10^-3^ | 1.00 |
| CX3CL1 | -0.12 | 9.52×10^-3^ | 1.00 |
| CDCP1 | 0.14 | 9.89×10^-3^ | 1.00 |
| MMP2 | -0.06 | 0.01 | 1.00 |
| ErbB3/HER3 | -0.11 | 0.01 | 1.00 |
| Nectin-4 | -0.12 | 0.01 | 1.00 |
| IL-18R1 | 0.13 | 0.01 | 1.00 |
| FGF-21 | 0.35 | 0.01 | 1.00 |
| Dkk-1 | 0.12 | 0.02 | 1.00 |
| FasL | -0.12 | 0.02 | 1.00 |
| FGF-19 | -0.17 | 0.02 | 1.00 |
| MIP-1 alpha | 0.13 | 0.02 | 1.00 |
| SLAMF1 | 0.11 | 0.02 | 1.00 |
| CXCL10 | 0.17 | 0.03 | 1.00 |
| TLR1 | 0.08 | 0.03 | 1.00 |
| EphB6 | -0.08 | 0.03 | 1.00 |
| Lymphotactin | 0.12 | 0.03 | 1.00 |
| CCL25 | -0.11 | 0.03 | 1.00 |
| SDF-1 | -0.09 | 0.03 | 1.00 |
| CCL3 | 0.11 | 0.03 | 1.00 |
| CEA | -0.08 | 0.03 | 1.00 |
| U-PAR | 0.11 | 0.03 | 1.00 |
| CD40L | 0.17 | 0.03 | 1.00 |
| CA125 | 0.16 | 0.03 | 1.00 |
| CT-like protein | -0.10 | 0.04 | 1.00 |
| TNFSF14 | 0.14 | 0.04 | 1.00 |
| HB-EGF | 0.12 | 0.04 | 1.00 |
| Amnionless | -0.12 | 0.04 | 1.00 |
| SIRP alpha | -0.10 | 0.04 | 1.00 |
| SPON1 | 0.12 | 0.05 | 1.00 |
| a-N-acetylgalactosaminidase | -0.09 | 0.05 | 1.00 |
| IL-13 R alpha 1 | -0.06 | 0.06 | 1.00 |
| E4-BP1 | 0.21 | 0.06 | 1.00 |
| BDNF | 0.62 | 0.06 | 1.00 |
| PAPP-A | 0.10 | 0.06 | 1.00 |
| Secretory carrier membrane protein 3 | 0.15 | 0.06 | 1.00 |
| ErbB4/HER4 | -0.08 | 0.07 | 1.00 |
| TLR3 | -0.11 | 0.07 | 1.00 |
| CD32 | -0.10 | 0.07 | 1.00 |
| MERTK | -0.12 | 0.08 | 1.00 |
| CD-6 | -0.10 | 0.08 | 1.00 |
| IL-1R-2 | -0.11 | 0.08 | 1.00 |
| PD-L1 | 0.07 | 0.08 | 1.00 |
| IFN-gamma-R2 | 0.13 | 0.08 | 1.00 |
| TNFRSF9 | 0.09 | 0.08 | 1.00 |
| MST1 | 0.12 | 0.08 | 1.00 |
| HMGB1 | 0.45 | 0.08 | 1.00 |
| MK | 0.10 | 0.09 | 1.00 |
| PD-L2 | -0.08 | 0.09 | 1.00 |
| Gastic intrinsic factor | -0.17 | 0.09 | 1.00 |
| NT-pro-BNP | 0.16 | 0.09 | 1.00 |
| IL-17RB | 0.10 | 0.09 | 1.00 |
| Gastrokine 1 | -0.08 | 0.09 | 1.00 |
| REN | 0.12 | 0.10 | 1.00 |
| PIgR | -0.07 | 0.10 | 1.00 |
| VEGF-D | -0.08 | 0.10 | 1.00 |
| TACE | 0.45 | 0.10 | 1.00 |
| LOX-1 | 0.13 | 0.11 | 1.00 |
| FGF-5 | -0.05 | 0.12 | 1.00 |
| ICAM-4 | -0.09 | 0.12 | 1.00 |
| ITLN-1 | -0.09 | 0.12 | 1.00 |
| IGF-I receptor | 0.07 | 0.12 | 1.00 |
| CD160 | -0.07 | 0.13 | 1.00 |
| CCL28 | 0.08 | 0.13 | 1.00 |
| TFPI-2 | 0.07 | 0.13 | 1.00 |
| Galectin-2 | -0.06 | 0.15 | 1.00 |
| ANXA1 | 0.10 | 0.16 | 1.00 |
| CASP-8 | -0.17 | 0.16 | 1.00 |
| Glypican-1 | -0.10 | 0.16 | 1.00 |
| FAS | -0.06 | 0.17 | 1.00 |
| GH | 0.26 | 0.17 | 1.00 |
| LAMP1 | -0.04 | 0.18 | 1.00 |
| GDNF | 0.05 | 0.19 | 1.00 |
| IL-17 RA | -0.07 | 0.19 | 1.00 |
| CD357 | -0.02 | 0.19 | 1.00 |
| TREML2 | -0.02 | 0.20 | 1.00 |
| CD4 | 0.05 | 0.20 | 1.00 |
| PAR-1 | -0.04 | 0.20 | 1.00 |
| Lymphotoxin beta R | 0.05 | 0.20 | 1.00 |
| IL-1R-1 | 0.16 | 0.21 | 1.00 |
| alpha-L-Iduronidase | 0.09 | 0.22 | 1.00 |
| Neurofibromin | 0.27 | 0.22 | 1.00 |
| STAMPB | -0.08 | 0.22 | 1.00 |
| MGMT | 0.09 | 0.23 | 1.00 |
| Dystroglycan | -0.08 | 0.23 | 1.00 |
| Notch 1 | 0.05 | 0.23 | 1.00 |
| CST5 | -0.05 | 0.24 | 1.00 |
| AXIN1 | 0.08 | 0.24 | 1.00 |
| ESM-1 | 0.07 | 0.24 | 1.00 |
| RANK | 0.05 | 0.24 | 1.00 |
| DC-SIGN | -0.03 | 0.25 | 1.00 |
| TYRO11 | -0.07 | 0.25 | 1.00 |
| TNFRSF14 | 0.15 | 0.25 | 1.00 |
| FS | 0.09 | 0.26 | 1.00 |
| CD244 | -0.05 | 0.26 | 1.00 |
| LAP TGF-beta | 0.10 | 0.26 | 1.00 |
| MCP-2 | -0.06 | 0.27 | 1.00 |
| Calgizzarin | 0.07 | 0.28 | 1.00 |
| Carbonic anhydrase 5A | -0.09 | 0.29 | 1.00 |
| PARK7 | 0.10 | 0.31 | 1.00 |
| LRP-6 | -0.04 | 0.31 | 1.00 |
| Galectin-9 | 0.08 | 0.31 | 1.00 |
| L-RAP | -0.04 | 0.32 | 1.00 |
| IL-3 R alpha | -0.05 | 0.33 | 1.00 |
| ST1A1 | -0.08 | 0.33 | 1.00 |
| TIE-2 | -0.05 | 0.34 | 1.00 |
| uPA | -0.04 | 0.34 | 1.00 |
| EN-RAGE | 0.07 | 0.37 | 1.00 |
| IL-12 | 0.06 | 0.37 | 1.00 |
| VEGFR-2 | -0.05 | 0.37 | 1.00 |
| TIM | -0.06 | 0.37 | 1.00 |
| MCP-1 | -0.05 | 0.38 | 1.00 |
| Hexosaminidase A/HEXA | 0.10 | 0.39 | 1.00 |
| ATR-interacting protein | 0.04 | 0.39 | 1.00 |
| PRSS8 | -0.05 | 0.39 | 1.00 |
| SCARB2 | -0.03 | 0.39 | 1.00 |
| SIRT2 | 0.07 | 0.40 | 1.00 |
| CD201 | -0.05 | 0.40 | 1.00 |
| CAIX | 0.06 | 0.40 | 1.00 |
| VIM | 0.05 | 0.40 | 1.00 |
| Brorin | -0.06 | 0.42 | 1.00 |
| R-spondin 3 | 0.04 | 0.42 | 1.00 |
| Superoxide dismutase | 0.07 | 0.42 | 1.00 |
| MIC-A | 0.09 | 0.43 | 1.00 |
| FADD | -0.06 | 0.43 | 1.00 |
| Peroxiredoxin 5 | 0.10 | 0.44 | 1.00 |
| PRL | 0.07 | 0.45 | 1.00 |
| N-Cadherin | -0.02 | 0.45 | 1.00 |
| Serine protease 27 | -0.03 | 0.45 | 1.00 |
| TWEAK | -0.03 | 0.46 | 1.00 |
| ADA | 0.04 | 0.47 | 1.00 |
| 2,4-dienoyl-CoA reductase | -0.05 | 0.48 | 1.00 |
| CD69 | -0.05 | 0.48 | 1.00 |
| Hydroxyacid oxidase 1 | -0.09 | 0.48 | 1.00 |
| Forkhead box protein O1 | 0.03 | 0.48 | 1.00 |
| NTPDase 6 | -0.03 | 0.49 | 1.00 |
| SLAMF5 | -0.02 | 0.49 | 1.00 |
| HSP-27 | 0.05 | 0.51 | 1.00 |
| CD23 | -0.04 | 0.52 | 1.00 |
| TM | -0.03 | 0.52 | 1.00 |
| Glutaredoxin 1 | -0.05 | 0.52 | 1.00 |
| CD27 Ligand | 0.03 | 0.52 | 1.00 |
| IL-12B | -0.04 | 0.54 | 1.00 |
| SLAMF2 | -0.03 | 0.54 | 1.00 |
| CG alpha | -0.04 | 0.54 | 1.00 |
| CD7 | 0.03 | 0.55 | 1.00 |
| PIGF | 0.02 | 0.58 | 1.00 |
| FLRG | 0.02 | 0.58 | 1.00 |
| IFN-gamma R1 | -0.02 | 0.59 | 1.00 |
| Integrin beta 2 | -0.02 | 0.59 | 1.00 |
| IL-16 | 0.03 | 0.63 | 1.00 |
| Cadherin-13 | 0.03 | 0.63 | 1.00 |
| THPO | -0.02 | 0.68 | 1.00 |
| Transglutaminase 2 | -0.03 | 0.68 | 1.00 |
| Granulysin | 0.02 | 0.69 | 1.00 |
| WIF1 | -0.02 | 0.69 | 1.00 |
| HO-1 | -0.05 | 0.69 | 1.00 |
| SLAMF3 | 0.02 | 0.70 | 1.00 |
| LYN | 0.01 | 0.72 | 1.00 |
| Dectin-1 | -0.02 | 0.73 | 1.00 |
| NK-p46 | 0.01 | 0.75 | 1.00 |
| eIF-4B | -0.03 | 0.75 | 1.00 |
| TNFRSF4 | 0.01 | 0.76 | 1.00 |
| CD90 | -0.03 | 0.77 | 1.00 |
| Max-like protein X | 0.01 | 0.78 | 1.00 |
| DAN | 0.02 | 0.78 | 1.00 |
| p27/Kip1/CDKN1B | -0.02 | 0.79 | 1.00 |
| EZR | 0.01 | 0.79 | 1.00 |
| CD40 | -0.01 | 0.79 | 1.00 |
| p70 S6 kinase alpha | 0.01 | 0.80 | 1.00 |
| EGF | -0.02 | 0.81 | 1.00 |
| GARP | -0.01 | 0.81 | 1.00 |
| FGF-23 | 0.01 | 0.81 | 1.00 |
| FSTL1 | -0.005 | 0.82 | 1.00 |
| TRAIL-R2 | -0.01 | 0.82 | 1.00 |
| MARCO | 0.01 | 0.84 | 1.00 |
| CD5 | -0.01 | 0.86 | 1.00 |
| Cathepsin H | 0.01 | 0.86 | 1.00 |
| LIF-R | 0.01 | 0.86 | 1.00 |
| MMP-7 | 0.01 | 0.88 | 1.00 |
| AGRP | 0.01 | 0.89 | 1.00 |
| GAL | 0.01 | 0.89 | 1.00 |
| Galectin-4 | 0.01 | 0.91 | 1.00 |
| PD-1 | 0.01 | 0.91 | 1.00 |
| VE-statin | 0 | 0.93 | 1.00 |
| TRAIL | 0 | 0.94 | 1.00 |
| HE4 | 0 | 0.94 | 1.00 |
| GITRL | 0 | 0.96 | 1.00 |
| CDKN1A | 0 | 0.96 | 1.00 |
| TACI | 0 | 0.96 | 1.00 |
| Protein Wnt-4 | 0 | 0.97 | 1.00 |
| beta-NGF | 0 | 0.97 | 1.00 |
| Sphingomyelin phosphodiesterase 3 | 0 | 0.97 | 1.00 |
| CTSL1 | 0 | 0.98 | 1.00 |
| BTLA | 0 | 0.98 | 1.00 |
| IL-10RB | 0 | 0.99 | 1.00 |
| MRAS | 0 | 0.99 | 1.00 |
| TFF3 | 0 | 1.00 | 1.00 |

**Supplementary Table 4:** Table of differentially expressed protein markers between Crohn’s disease (CD) cases and controls in serum with age and sex as covariates. Log_2_ Fold-Change denotes mean CD expression – mean control expression, with expression as normalised Ct values.

| Protein | Log_2_FC | P Value | Holm P |
| --- | --- | --- | --- |
| CXCL9 | 1.02 | 1.60×10^-17^ | 5.00×10^-15^ |
| OSM | 0.82 | 1.84×10^-14^ | 5.75×10^-12^ |
| MMP-12 | 0.65 | 1.47×10^-13^ | 4.59×10^-11^ |
| HGF | 0.52 | 4.95×10^-12^ | 1.54×10^-9^ |
| CXCL11 | 0.59 | 7.09×10^-11^ | 2.19×10^-8^ |
| IL-6 | 0.87 | 1.30×10^-10^ | 4.01×10^-8^ |
| IL-8 | 0.61 | 1.49×10^-10^ | 4.58×10^-8^ |
| IFN-gamma | 0.65 | 1.73×10^-10^ | 5.30×10^-8^ |
| CXCL1 | 0.58 | 3.12×10^-10^ | 9.52×10^-8^ |
| Gas6 | -0.22 | 3.57×10^-10^ | 1.09×10^-7^ |
| CCL20 | 0.63 | 2.68×10^-9^ | 8.11×10^-7^ |
| ITGAV | -0.28 | 4.80×10^-9^ | 1.45×10^-6^ |
| CXCL2/3 | 0.76 | 5.21×10^-9^ | 1.57×10^-6^ |
| IL-17A | 0.44 | 5.75×10^-9^ | 1.73×10^-6^ |
| gp130 | -0.33 | 6.26×10^-9^ | 1.87×10^-6^ |
| FREB-2 | 0.34 | 1.80×10^-8^ | 5.36×10^-6^ |
| CSF-1 | 0.40 | 6.00×10^-8^ | 1.78×10^-5^ |
| VEGF-A | 0.43 | 1.44×10^-7^ | 4.26×10^-5^ |
| IL-1Ra | 0.49 | 1.60×10^-7^ | 4.71×10^-5^ |
| LITAF | 0.54 | 2.18×10^-7^ | 6.42×10^-5^ |
| SCF | -0.41 | 1.44×10^-6^ | 4.23×10^-4^ |
| IL-4 R alpha | 0.30 | 1.85×10^-6^ | 5.42×10^-4^ |
| Ep-CAM | -0.50 | 2.35×10^-6^ | 6.84×10^-4^ |
| Decoy receptor 3 | 0.59 | 2.42×10^-6^ | 7.01×10^-4^ |
| BOC | -0.21 | 3.83×10^-6^ | 1.11×10^-3^ |
| IL-17C | 0.31 | 4.88×10^-6^ | 1.41×10^-3^ |
| FR-alpha | -0.21 | 8.98×10^-6^ | 2.58×10^-3^ |
| MMP-1 | 0.50 | 1.04×10^-5^ | 2.96×10^-3^ |
| IL-7 | 0.30 | 1.11×10^-5^ | 3.15×10^-3^ |
| CXCL5 | 0.41 | 1.22×10^-5^ | 3.47×10^-3^ |
| ICOSLG | -0.40 | 1.25×10^-5^ | 3.53×10^-3^ |
| TRAIL R1 | 0.22 | 1.47×10^-5^ | 4.15×10^-3^ |
| DNER | -0.25 | 1.58×10^-5^ | 4.45×10^-3^ |
| CCL11 | 0.27 | 1.72×10^-5^ | 4.81×10^-3^ |
| CDH3 | -0.24 | 2.06×10^-5^ | 5.76×10^-3^ |
| PTPN22 | 0.42 | 2.88×10^-5^ | 8.01×10^-3^ |
| REG-4 | 0.35 | 4.42×10^-5^ | 1.22×10^-2^ |
| Carboxypeptidase M | -0.17 | 4.84×10^-5^ | 1.34×10^-2^ |
| FUR | 0.24 | 5.33×10^-5^ | 1.47×10^-2^ |
| TRANCE | -0.35 | 5.74×10^-5^ | 1.57×10^-2^ |
| ITGA1 | -0.25 | 6.78×10^-5^ | 1.85×10^-2^ |
| CCL23 | 0.32 | 7.03×10^-5^ | 1.91×10^-2^ |
| TREM-1 | 0.21 | 8.12×10^-5^ | 2.20×10^-2^ |
| MCP-4 | -0.25 | 9.09×10^-5^ | 2.46×10^-2^ |
| NOV | -0.14 | 9.41×10^-5^ | 2.53×10^-2^ |
| TLR2 | 0.19 | 1.02×10^-4^ | 2.74×10^-2^ |
| BAFF | 0.21 | 1.09×10^-4^ | 2.90×10^-2^ |
| MCP-3 | 0.27 | 1.12×10^-4^ | 2.97×10^-2^ |
| FGF-19 | -0.34 | 1.22×10^-4^ | 3.24×10^-2^ |
| ILT-3 | 0.20 | 1.41×10^-4^ | 3.72×10^-2^ |
| NT-3 | -0.18 | 1.42×10^-4^ | 3.72×10^-2^ |
| TGFA | 0.35 | 1.46×10^-4^ | 3.83×10^-2^ |
| TNFB | -0.23 | 1.50×10^-4^ | 3.92×10^-2^ |
| CD89 | 0.26 | 1.58×10^-4^ | 4.10×10^-2^ |
| TNF-R1 | 0.24 | 1.62×10^-4^ | 4.18×10^-2^ |

**Supplementary Table 5:** Table of differentially expressed protein markers between Ulcerative Colitis (UC) cases and controls in serum with age and sex as covariates. Log_2_ Fold-Change denotes mean UC expression – mean control expression, with expression as normalised Ct values.

| Protein | Log_2_FC | P Value | Holm P |
| --- | --- | --- | --- |
| MMP-12 | 1.14 | 1.15×10^-28^ | 3.59×10^-26^ |
| Granzyme B | 1.54 | 2.54×10^-25^ | 7.92×10^-23^ |
| IL-17A | 0.95 | 1.51×10^-23^ | 4.69×10^-21^ |
| IL-8 | 1.08 | 1.28×10^-22^ | 3.96×10^-20^ |
| MMP-10 | 0.97 | 1.94×10^-20^ | 6.01×10^-18^ |
| CXCL1 | 0.96 | 1.13×10^-19^ | 3.48×10^-17^ |
| AR | 0.71 | 2.16×10^-16^ | 6.62×10^-14^ |
| OSM | 0.85 | 5.51×10^-14^ | 1.69×10^-11^ |
| IL-17C | 0.49 | 5.23×10^-11^ | 1.59×10^-8^ |
| CXCL9 | 0.72 | 5.50×10^-10^ | 1.67×10^-7^ |
| CXCL11 | 0.60 | 6.52×10^-10^ | 1.98×10^-7^ |
| TRANCE | -0.57 | 6.59×10^-10^ | 1.99×10^-7^ |
| CCL20 | 0.66 | 1.06×10^-9^ | 3.18×10^-7^ |
| CXCL2/3 | 0.80 | 2.60×10^-9^ | 7.81×10^-7^ |
| IL-10 | 0.67 | 2.86×10^-9^ | 8.54×10^-7^ |
| HGF | 0.48 | 5.76×10^-9^ | 1.72×10^-6^ |
| TGFA | 0.52 | 1.77×10^-8^ | 5.27×10^-6^ |
| ITGAV | -0.27 | 2.13×10^-8^ | 6.30×10^-6^ |
| FR-alpha | -0.26 | 2.57×10^-8^ | 7.59×10^-6^ |
| Decoy receptor 3 | 0.66 | 2.79×10^-8^ | 8.21×10^-6^ |
| Langerin | -0.29 | 2.93×10^-8^ | 8.60×10^-6^ |
| NOV | -0.20 | 6.46×10^-8^ | 1.89×10^-5^ |
| IL-6 | 0.81 | 7.76×10^-8^ | 2.26×10^-5^ |
| IL-17D | -0.19 | 1.28×10^-7^ | 3.70×10^-5^ |
| Galectin-1 | -0.19 | 1.43×10^-7^ | 4.12×10^-5^ |
| Carboxypeptidase M | -0.21 | 2.42×10^-7^ | 6.97×10^-5^ |
| TROY | -0.23 | 3.11×10^-7^ | 8.92×10^-5^ |
| IL-1Ra | 0.48 | 5.51×10^-7^ | 1.58×10^-4^ |
| IL-4 R alpha | 0.31 | 8.01×10^-7^ | 2.28×10^-4^ |
| Gas6 | -0.16 | 8.48×10^-7^ | 2.41×10^-4^ |
| GFR alpha-3 | -0.21 | 1.52×10^-6^ | 4.29×10^-4^ |
| CCL11 | 0.31 | 1.91×10^-6^ | 5.39×10^-4^ |
| SCF | -0.40 | 2.40×10^-6^ | 6.75×10^-4^ |
| Ret | -0.25 | 4.02×10^-6^ | 1.13×10^-3^ |
| REG-4 | 0.40 | 4.50×10^-6^ | 1.25×10^-3^ |
| PTPN22 | 0.45 | 9.88×10^-6^ | 2.75×10^-3^ |
| Gastrotropin | -0.24 | 1.02×10^-5^ | 2.81×10^-3^ |
| PSGL-1 | -0.13 | 1.45×10^-5^ | 4.00×10^-3^ |
| Leukocyte tyrosine kinase receptor | -0.13 | 1.59×10^-5^ | 4.36×10^-3^ |
| EDIL3 | -0.26 | 1.59×10^-5^ | 4.36×10^-3^ |
| ICOSLG | -0.40 | 1.78×10^-5^ | 4.87×10^-3^ |
| BMP-9 | -0.30 | 4.12×10^-5^ | 1.12×10^-2^ |
| ErbB2/HER2 | -0.24 | 5.19×10^-5^ | 1.41×10^-2^ |
| DNER | -0.25 | 5.33×10^-5^ | 1.44×10^-2^ |
| RAGE | -0.24 | 1.10×10^-4^ | 2.96×10^-2^ |
| IL-27 | 0.21 | 1.60×10^-4^ | 4.30×10^-2^ |

**Supplementary Table 6:** Table of differentially expressed protein markers between Ulcerative Colitis (UC) cases and Crohn’s disease (CD) in serum with age and sex as covariates. Log_2_ Fold-Change denotes mean UC expression – mean CD expression, with expression as normalised Ct values.

| Protein | Log_2_FC | P Value | Holm P |
| --- | --- | --- | --- |
| Granzyme B | -1.13 | 3.92×10^-8^ | 1.23×10^-5^ |
| MMP-10 | -0.65 | 4.87×10^-7^ | 1.52×10^-4^ |
| AR | -0.47 | 7.98×10^-6^ | 2.48×10^-3^ |
| MMP-12 | -0.51 | 1.50×10^-5^ | 4.66×10^-3^ |
| IL-17A | -0.53 | 1.63×10^-5^ | 5.02×10^-3^ |

**Supplementary Table 7:** Disease outcomes in patients with Ulcerative colitis (UC) and Crohn’s disease (CD).

|  | Treatment escalation  N=67 | No treatment escalation  N=212 |
| --- | --- | --- |
| Censored follow-up | | |
| Median (IQR) | 116 (39.5-236.5) | 583 (439.8-853.0) |
| Primary Treatment prior to escalation | | |
| 5ASA (+Steroid) | 14 (7) | 129 (21) |
| 6MP (+Steroid) | 7 (0) | 0 (0) |
| AZA (+Steroid) | 28 (7) | 16 (12) |
| Steroid (IV) | 12 (3) | 26 (0) |
| Ciclosporin (+IV Steroids) | 4 (4) | 1 (0) |
| Anti-TNF (+Other) | 0 | 6 (5) |
| Other | 2 | 2 |
| None | 0 | 32 |
| Secondary Treatment at escalation | | |
| Anti-TNF | 43 |  |
| Anti-TNF + Other | 6 |  |
| Ciclosporin | 3 |  |
| Surgery | 15 |  |

### Footnote: AZA: azathioprine; 5ASA: aminosalicylates; 6MP: mercaptopurine.

5ASA (+Steroid) value 14 (7) denotes 14 patients on 5ASA of whom 7 were also on steroids.

**Supplemental Table 8**: Proteins associated with treatment escalation (anti-TNF/ciclosporin and/or surgery) in UC patients. HR: hazard ratio associated with a one unit increase in log_2_ expression, IQR HR: hazard ratio associated with moving between the 25^th^ and 75^th^ percentile of expression in the direction of increased risk.

| **Protein** | **P value** | **HR** | **IQR HR** | **Holm P value** |
| --- | --- | --- | --- | --- |
| **IL-1RA** | 3.48×10^-7^ | 2.38 | 3.74 | 1.09×10^-4^ |
| **CSF-1** | 3.66×10^-7^ | 3.63 | 3.69 | 1.14×10^-4^ |
| **TNF-R1** | 2.33×10^-6^ | 5.00 | 4.58 | 7.26×10^-4^ |
| **ITGAV** | 2.86×10^-6^ | 0.13 | 10.53 | 8.85×10^-4^ |
| **IL-8** | 3.67×10^-6^ | 2.04 | 3.34 | 1.13×10^-3^ |
| **AM** | 5.44×10^-6^ | 3.64 | 3.81 | 1.68×10^-3^ |
| **CD6** | 9.35×10^-6^ | 0.20 | 5.85 | 2.87×10^-3^ |
| **LTBR** | 1.04×10^-5^ | 7.99 | 5.20 | 3.18×10^-3^ |
| **CCL23** | 1.25×10^-5^ | 3.15 | 3.75 | 3.82×10^-3^ |
| **EpCAM** | 1.51×10^-5^ | 0.41 | 1.68 | 4.60×10^-3^ |
| **IL-17A** | 1.54×10^-5^ | 1.83 | 2.89 | 4.68×10^-3^ |
| **IL-18** | 2.43×10^-5^ | 3.07 | 3.23 | 7.34×10^-3^ |
| **MMP-10** | 2.98×10^-5^ | 1.77 | 2.37 | 8.97×10^-3^ |
| **MMP-12** | 3.10×10^-5^ | 2.58 | 4.33 | 9.31×10^-3^ |
| **FasL** | 3.35×10^-5^ | 0.26 | 5.60 | 0.010 |
| **TRAILR1** | 5.57×10^-5^ | 2.82 | 2.12 | 0.017 |
| **TRANCE** | 7.90×10^-5^ | 0.42 | 1.93 | 0.023 |
| **RANK** | 8.14×10^-5^ | 6.01 | 3.44 | 0.024 |
| **OPG** | 9.29×10^-5^ | 3.05 | 2.51 | 0.027 |
| **HGF** | 1.27×10^-4^ | 1.80 | 2.02 | 0.037 |
| **FRalpha** | 1.59×10^-4^ | 0.14 | 10.69 | 0.047 |
| **TYRO11** | 1.67×10^-4^ | 16.63 | 5.72 | 0.049 |

**Supplementary Table 9**: All significant pQTLs in proteins which had significant IBD-associated expression changes.

| Gene | SNP | P Value | Holm P | AA | Aa | aa |
| --- | --- | --- | --- | --- | --- | --- |
| VEGF-A | rs7767396 | 9.2×10^-22^ | 8.7×10^-18^ | 164 | 264 | 121 |
| NOV | rs2071518 | 2.6×10^-19^ | 2.4×10^-15^ | 283 | 227 | 39 |
| CXCL5 | rs425535 | 2.9×10^-19^ | 2.7×10^-15^ | 442 | 96 | 11 |
| NOV | rs1461693 | 3.4×10^-18^ | 3.2×10^-14^ | 297 | 217 | 35 |
| VEGF-A | rs9472159 | 1.7×10^-17^ | 1.6×10^-13^ | 148 | 268 | 132 |
| CXCL5 | rs352010 | 2.4×10^-17^ | 2.3×10^-13^ | 441 | 97 | 11 |
| NOV | rs2071519 | 3.5×10^-16^ | 3.3×10^-12^ | 304 | 211 | 34 |
| NOV | rs4871221 | 4.9×10^-16^ | 4.7×10^-12^ | 368 | 161 | 20 |
| VEGF-A | rs9472158 | 5.7×10^-15^ | 5.3×10^-11^ | 191 | 258 | 100 |
| VEGF-A | rs9369434 | 7.9×10^-15^ | 7.5×10^-11^ | 190 | 258 | 101 |
| FREB-2 | rs2333749 | 3.3×10^-13^ | 3.1×10^-9^ | 227 | 250 | 70 |
| CXCL5 | rs352024 | 6.2×10^-12^ | 5.9×10^-8^ | 436 | 103 | 10 |
| CXCL5 | rs352018 | 6.2×10^-12^ | 5.9×10^-8^ | 436 | 103 | 10 |
| CXCL5 | rs487834 | 6.2×10^-12^ | 5.9×10^-8^ | 436 | 103 | 10 |
| MMP-1 | rs479095 | 2.6×10^-11^ | 2.4×10^-7^ | 297 | 215 | 37 |
| LTK | rs1200353 | 4.7×10^-11^ | 4.4×10^-7^ | 268 | 232 | 49 |
| VEGF-A | rs9472173 | 5.8×10^-11^ | 5.4×10^-7^ | 146 | 274 | 129 |
| CXCL5 | rs4694656 | 6.2×10^-11^ | 5.8×10^-7^ | 429 | 110 | 10 |
| NOV | rs17187503 | 6.8×10^-11^ | 6.5×10^-7^ | 330 | 194 | 25 |
| NOV | rs17187655 | 9.9×10^-11^ | 9.3×10^-7^ | 331 | 193 | 25 |
| NOV | rs11785549 | 9.9×10^-11^ | 9.3×10^-7^ | 331 | 193 | 25 |
| NOV | rs11526 | 9.9×10^-11^ | 9.3×10^-7^ | 331 | 193 | 25 |
| CXCL5 | rs4694180 | 1.1×10^-10^ | 9.9×10^-7^ | 372 | 153 | 24 |
| CXCL5 | rs10938105 | 1.3×10^-10^ | 1.3×10^-6^ | 370 | 155 | 24 |
| LTK | rs1473781 | 1.4×10^-10^ | 1.3×10^-6^ | 229 | 253 | 67 |
| CXCL5 | rs13122604 | 1.9×10^-10^ | 1.8×10^-6^ | 386 | 145 | 18 |
| CXCL5 | rs183028 | 2.4×10^-10^ | 2.3×10^-6^ | 444 | 91 | 12 |
| MMP-1 | rs1010698 | 3.8×10^-10^ | 3.6×10^-6^ | 142 | 265 | 141 |
| CXCL5 | rs4694655 | 3.8×10^-10^ | 3.6×10^-6^ | 361 | 162 | 26 |
| MMP-1 | rs11225434 | 4.2×10^-10^ | 3.9×10^-6^ | 157 | 259 | 133 |
| CXCL5 | rs2472649 | 7.3×10^-10^ | 6.9×10^-6^ | 390 | 138 | 20 |
| CCL23 | rs860559 | 7.4×10^-10^ | 7.0×10^-6^ | 341 | 172 | 36 |
| MMP-1 | rs519806 | 7.5×10^-10^ | 7.0×10^-6^ | 192 | 256 | 100 |
| CCL23 | exm1313428 | 1.1×10^-9^ | 1.0×10^-5^ | 343 | 170 | 36 |
| CCL23 | rs1617208 | 1.2×10^-9^ | 1.1×10^-5^ | 344 | 169 | 36 |
| VEGF-A | rs729391 | 3.9×10^-9^ | 3.7×10^-5^ | 215 | 247 | 86 |
| CXCL5 | rs1595009 | 1.0×10^-8^ | 9.8×10^-5^ | 367 | 155 | 26 |
| CCL23 | rs6505503 | 1.1×10^-8^ | 0.00011 | 358 | 162 | 29 |
| MMP-1 | rs645419 | 1.4×10^-8^ | 0.00013 | 144 | 264 | 140 |
| MMP-1 | exm951505 | 2.3×10^-8^ | 0.00022 | 143 | 264 | 140 |
| LTK | rs6493004 | 3.1×10^-8^ | 3.0×10^-4^ | 273 | 230 | 46 |
| CD89 | rs1865096 | 9.4×10^-8^ | 0.00089 | 289 | 205 | 55 |
| LTK | exm1151064 | 1.6×10^-7^ | 0.0015 | 441 | 100 | 8 |
| LTK | rs2305030 | 1.6×10^-7^ | 0.0015 | 441 | 100 | 8 |
| CCL23 | rs854692 | 2.8×10^-7^ | 0.0027 | 324 | 185 | 40 |
| LTK | rs9944249 | 4.5×10^-7^ | 0.0043 | 183 | 280 | 86 |
| NOV | rs1058913 | 7.8×10^-7^ | 0.0073 | 326 | 196 | 27 |
| MMP-12 | rs17368814 | 7.8×10^-7^ | 0.0073 | 438 | 102 | 9 |
| MMP-1 | rs495366 | 9.6×10^-7^ | 0.009 | 296 | 215 | 37 |
| MMP-1 | rs522616 | 1.1×10^-6^ | 0.01 | 345 | 180 | 22 |
| FREB-2 | rs12135523 | 1.2×10^-6^ | 0.011 | 139 | 293 | 117 |
| FREB-2 | rs7529425 | 1.5×10^-6^ | 0.014 | 429 | 117 | 3 |
| MMP-12 | exm951532 | 1.6×10^-6^ | 0.015 | 439 | 101 | 9 |
| NOV | rs16892729 | 1.6×10^-6^ | 0.015 | 320 | 197 | 31 |
| MMP-1 | rs1144397 | 1.6×10^-6^ | 0.016 | 191 | 264 | 94 |
| FREB-2 | rs6427618 | 2.2×10^-6^ | 0.02 | 180 | 285 | 84 |
| LTK | rs1918310 | 3.0×10^-6^ | 0.028 | 244 | 245 | 60 |
| VEGF-A | rs7764227 | 3.6×10^-6^ | 0.034 | 288 | 209 | 52 |
| LTK | exm1151599 | 4.5×10^-6^ | 0.042 | 235 | 249 | 65 |

**References**

1	Assarsson E, Lundberg M, Holmquist G, *et al.* Homogenous 96-plex PEA immunoassay exhibiting high sensitivity, specificity, and excellent scalability. *PLoS One* 2014; **9**: e95192.

2	Lind L, Ärnlöv J, Lindahl B, Siegbahn A, Sundström J, Ingelsson E. Use of a proximity extension assay proteomics chip to discover new biomarkers for human atherosclerosis. *Atherosclerosis* 2015; **242**: 205–10.

3	Shabalin AA. Matrix eQTL: ultra fast eQTL analysis via large matrix operations. *Bioinformatics* 2012; **28**: 1353–8.
